# Local control of resource allocation is sufficient to model optimal dynamics in syntrophic systems

**DOI:** 10.1101/787465

**Authors:** Glenn Ledder, Sabrina E. Russo, Erik B. Muller, Angela Peace, Roger M. Nisbet

## Abstract

Syntrophic systems are common in nature and include forms of obligate mutualisms in which each participating organism or component of an organism obtains from the other an essential nutrient or metabolic product that it cannot provide for itself. Models of how these complementary resources are allocated between partners often assume optimal behavior, but whether mechanisms enabling global control exist in syntrophic systems, and what form they might take, is unknown. Recognizing that growth of plant organs that supply complementary resources, like roots and shoots, can occur autonomously, we present a theory of plant growth in which root-shoot allocation is determined by purely local rules. Each organ uses as much as it can of its locally produced or acquired resource (inorganic nitrogen or photosynthate) and shares only the surplus. Subject to stoichiometric conditions that likely hold for most plants, purely local rules produce the same optimal allocation as would global control, even in a fluctuating environment, with sharing the surplus being the specific mechanism stabilizing syntrophic dynamics. Our local control model contributes a novel approach to plant growth modeling because it assumes a simple mechanism of root:shoot allocation that can be considered a higher-level physiological rule, from which the optimal growth outcome emerges from the system’s dynamics, rather than being built into the model. Moreover, our model is general, in that the mechanism of sharing the surplus can readily be adapted to many obligate syntrophic relationships.

## 1 Introduction

Nature offers many examples of obligate mutualisms in which participating organisms exchange resources, such as energy, essential micro- or macro-nutrients, or metabolic products, that they cannot provide for themselves. The resulting partnership may take the form of a holobiont (Mindell 1992; Rohwer et al. 2002; Bordenstein & Theis 2015), such as (1) dinoflagellates coexisting with corals, jellyfish, or molluscs (Muscatine & Porter 1977), (2) an association of free-living organisms such as leaf-cutter ants and fungi (Kang et al, 2011), or (3) engineered non-mating yeast cells where each supplies an essential metabolite to the other strain (Shou et al 2007).

The allocation of complementary resources to organs within the same organism is analogous to resource-sharing between partners in a holobiont. An example comes from plants, which, unlike unitary animals, are modular, in that growth of organs such as roots and shoots can occur somewhat autonomously (Haukioja 1991). Roots and shoots supply complementary resources to the whole plant, which is also similar to resource-sharing in a holobiont: roots supply water and nutrients required by the shoots for photosynthesis and the construction of photosynthetic tissues, while the shoots synthesize carbohydrates that are required for cellular respiration and tissue construction in the root. How the plant’s resources are allocated to these organs is a principal determinant of whole-plant growth, survival, and reproduction, and, ultimately, the environment to which a plant species is adapted (Reich 2002; Poorter et al. 2012). Moreover, the ratio between root and shoot biomass plastically responds to the environment (Reich 2002; Weiner 2004), leading to the hypothesis that plants, like holobionts, allocate resources between these organs (or partners) in a way that maximizes fitness (Cody 1966; Harper & Ogden 1970). In contrast to organs within a plant, individuals interacting in a syntrophic holobiont have distinct genotypes. From an evolutionary perspective, natural selection should operate to produce ecological resource allocation strategies that independently maximize the fitness of each organism (Bordenstein & Theis 2015; Moran & Sloan 2015). The processes regulating how individuals (in the case of the holobiont) or organs (in the case of a single organism) share resources are poorly understood, but the diverse ecological contexts and the evolutionary persistence of these partnerships across a broad range of taxa suggest that general dynamical mechanisms may be involved. Dynamic models of the control of resource-sharing in such partnerships are therefore fundamental to understanding their functional ecology and evolution.

Many models describing resource allocation and growth in plants take an evolutionary approach, in which resources are distributed among competing functions in a way that maximizes a whole-plant proxy for fitness, such as whole-plant growth rate or reproductive output from an annual plant (Brouwer 1963, 1983; Bloom et al. 1985; Wilson 1988; Franklin et al. 2012). Drawing from economic theory, the functional equilibrium or balanced growth hypothesis states that endogenous resources are optimally allocated among competing processes so that each resource limits all processes to the same degree (Bloom et al. 1985). Applied to roots and shoots, the functional equilibrium hypothesis predicts that endogenous resources are preferentially and plastically partitioned to the above or belowground compartment that acquires the exogenous resource currently most limiting to whole plant growth in a changing environment (Bloom et al. 1985; Lerdau 1992), a strategy that has been observed in real plants (Weiner 2004; Poorter & Nagel 2000). Implicit in the use of dynamic optimization to model resource allocation and growth is that there is some form of “global” (e.g., hormonal) control at the level of the whole organism that can sense deficiency in a particular function and then adjust allocation in such a way as to achieve and maintain the optimal allocation. For example, Iwasa & Roughgarden (1984) used a plant model in which the availability of a single resource, photosynthate, determines plant growth, with the rate of its accumulation assumed to be a function of shoot and root biomasses. They assumed that the time-dependent proportions of photosynthate allocated to root, shoot, and fruit are controllable at the level of the whole organism, with allocation chosen so as to optimize reproduction over the lifetime of the plant. Velten & Richter (1995) studied in detail a simpler model in which the objective was to maximize vegetative biomass (root plus shoot). Nevertheless, the assumption of global control is implicit in all such models, else there would be no way to achieve the optimal solution.

Other models still assume an objective function that is maximized, but also specify some physiological detail as to how that optimality is achieved. Optimal allocation outcomes have been achieved using the Liebig minimum rule to determine how resources limit whole-plant growth and metabolic scaling rules to determine the rate of tissue production (Lin et al. 2014). Alternatively, many models assume total photosynthetic carbon gain is maximized and provide a physiological function that dictates how much photosynthate is gained, given particular biomasses of root and shoot (e.g., Sterck & Schieving 2011; Reynolds & Pacala 1992). There are, however, a few complications of assuming that there is a global controller for resource-allocation or sharing in an optimization setting (Wilson 1988; Cheeseman 1993). First, allocation patterns are the product of several interacting physiological processes, not one process (Cannell & Dewar 1994). Second, it is not known if such an “optimizing” global controller that could integrate these processes exists for single organisms or organisms involved in syntrophic relationships. Alternatively, for example, in plants natural selection may have operated to produce a set point for root:shoot biomass ratio for reasons that may be unrelated to optimality. When perturbed off the set point, the plant may grow in a way that reestablishes this genetically determined functional equilibrium (Reich 2002). Third, in terms of modeling, one needs to define a priori the quantity to be optimized and to provide mechanisms dictating how that optimality is achieved. Yet, the physiological mechanisms that control resource allocation are poorly understood, even in plants, for which models of root:shoot allocation have a long history (Wilson 1988).

An alternative to making ad hoc assumptions about an objective function for optimization is an ecologically based approach to resource allocation between roots and shoots that involves considering the interactions between a plant and its conspecifics in a population or community in a game-theoretical context (King 1993; McNickle and Dybzinski 2013). Farrior et al. (2013) used a physiological plant model and explicit competitive interactions for water in a finite habitat. This approach allowed calculation of evolutionarily stable strategies (ESSs) without the a priori choice of an objective function, as the ESS represents an allocation strategy that cannot be invaded by an alternative phenotype (Farrior et al. 2013), an approach also used to model above vs. belowground carbon allocation at the forest stand scale (Dybzinski et al. 2011). While still assuming global control, this approach does not require hypothesizing an arbitrary target for optimization, since the outcome is that a plant allocates resources to roots and shoots in whatever way makes it the most competitive in the community.

A separate body of literature starts at the level of physiology and does not explicitly consider evolutionary or competitive processes. Thornley (1972) proposed that root-shoot ratio could be regulated without global control, but through a representation of source, transport, and sink processes associated with differential resource capture and use by roots and shoots, which he termed the transport-resistance (TR) framework (Thornley 1998). A two-compartment, two-resource model, the TR framework assumes a mechanism for local control of resource distribution between organs: a translocation rule that promotes equalization of the concentrations of resources (carbon and nitrogen) in root and shoot and that is determined passively by differences in the concentrations and resistances to their translocation between organs. Effectively, translocation processes compete with the organs’ utilization processes for the resources, so that even when the uptake capacity of an organ is deficient, relative to what is required for maximal growth, some of the resource that it collects will still be transferred to the other organ. The TR framework has been incorporated in more elaborate physiological models, a recent example being Feller et al., (2015). Other models of plant growth and resource allocation to roots and shoots that do not include some form of global control have used strictly local rules to determine allocation (Cheeseman 1993; Cheeseman et al. 1996). With two compartments and two resources (carbon and nitrogen), the rule to distribute the resources was that each organ automatically supplies the other with a fixed percentage of the resource that it acquires. While this rule implies a local mechanism, it does not involve any form of control, since a fixed parameter controls the resource distribution, which thus cannot change if conditions change. The growth rates of root and shoot are determined by multi-parameter polynomial functions of the concentrations of carbon and nitrogen in the organs. Thus, the good fit to data of this model likely was achieved because of its many parameters, rather than because of the local control mechanism per se.

We offer a new approach to modeling plant growth and root:shoot allocation—the local control theory of plant resource allocation—that derives from recent theory for obligate syntrophic symbiosis. Our approach is an advance because it assumes a simple mechanism of root:shoot allocation, which we refer to as sharing the surplus, from which the optimal plant growth and allocation outcomes emerge from the systems dynamics, rather than being built into the model or being contingent upon the composition of the local competitive community. Instead of invoking specific physiological mechanisms controlling movement of resources between roots and shoots, such as translocation resistance, we assume that each operates selfishly, and our model is agnostic as to the specific physiological mechanisms involved. We draw on previous models on reef corals (Muller et al. 2009; Cunning et al. 2017), in which intracellular dinoflagellates of the genus Symbiodinium perform photosynthesis using nutrients acquired by the animal host, and the animal in turn uses photosynthate from the symbiont as a source of chemical energy and carbon. Host and symbiont were assumed to operate “selfishly,” making only surplus resource or metabolic product available to the partner. Such local control of resource sharing offers an alternative to global control and has the advantage of not requiring assumptions as to how global control operates. We constructed a conceptually similar ordinary differential model for a plant with inorganic nitrogen and photosynthate as the shared resources (Kooijman 2010). Roots and shoots are each able to supply only one of the resources, either through assimilation from the environment (roots and nitrogen, analogous to the coral) or through synthesis (shoots and photosynthate, analogous to *Symbiodinium*). There is no global control of resource sharing, as each partner only passes surplus to the other, in keeping with early hypotheses for the regulation of root and shoot growth (White 1937). Each partner’s biomass production utilizes the resources in a fixed stoichiometric ratio (Davidson 1969; Garnier 1991). Partners may differ in the extent to which each needs the resource supplied by the other, in keeping with stoichiometric differences between organs such as roots and shoots. Production kinetics are modeled as a function of the input streams of the two resources. Resource assimilation rates are modeled as a function of the relevant partner’s biomass (Muller et al. 2001).

We used linearized stability analysis to derive conditions for achieving a stable equilibrium between the assimilation capacities of roots and shoots. This analysis shows that the passive allocation system of relying on sharing of surplus resources as a mechanism of local control achieves the same growth rate and biomass allocation as could be achieved by a hypothetical global controller, under a broad range of conditions. An investigation of the transient dynamics that occur in response to changes in environmental conditions or a drastic loss of root or shoot tissue shows that the response achieved via local control is actually superior in many scenarios to the expectations of the functional equilibrium hypothesis described above.

## 2 A Model for Growth of an Idealized Plant

Our idealized plant has two components, called “roots” and “shoots”. Their biomasses, denoted by *R*(*t*) and *S*(*t*) respectively, are the primary state variables in our model. The former is an abstraction of the organ responsible for assimilating water and macronutrients, while the latter is an abstraction of the organ for absorbing light and assimilating carbon into photosynthate. We call the principal macronutrient “N” and the photosynthate “C”. The “biomass” of each component is defined to include only biologically-active tissues, and therefore does not include xylem, cork, or bark, which are functional tissues comprised of dead cells. Formation of functional tissues comprised of dead cells, for example, development of xylem from the vascular cambium, is considered to be a portion of the turnover of root and shoot biomasses.

We assume that assimilated resources are used immediately to create new root and shoot biomass, with no explicit incorporation of time delays or storage. While storage of carbohydrates and nutrients is known to occur in plants (Chapin et al. 1990), the dynamics, physiological control, and functional relevance of storage is debated (Sala et al. 2010), and our focus is on elucidating the *dynamical properties* of a resource allocation model that does not assume global control. We discuss the potential impact of reserves on dynamics in the Discussion.

The core assumptions, and the notation for state variables and for the flows of C and N, are shown in Figure 1 and listed below. (See Table 1 for a summary of notation.)

**Fig. 1.**
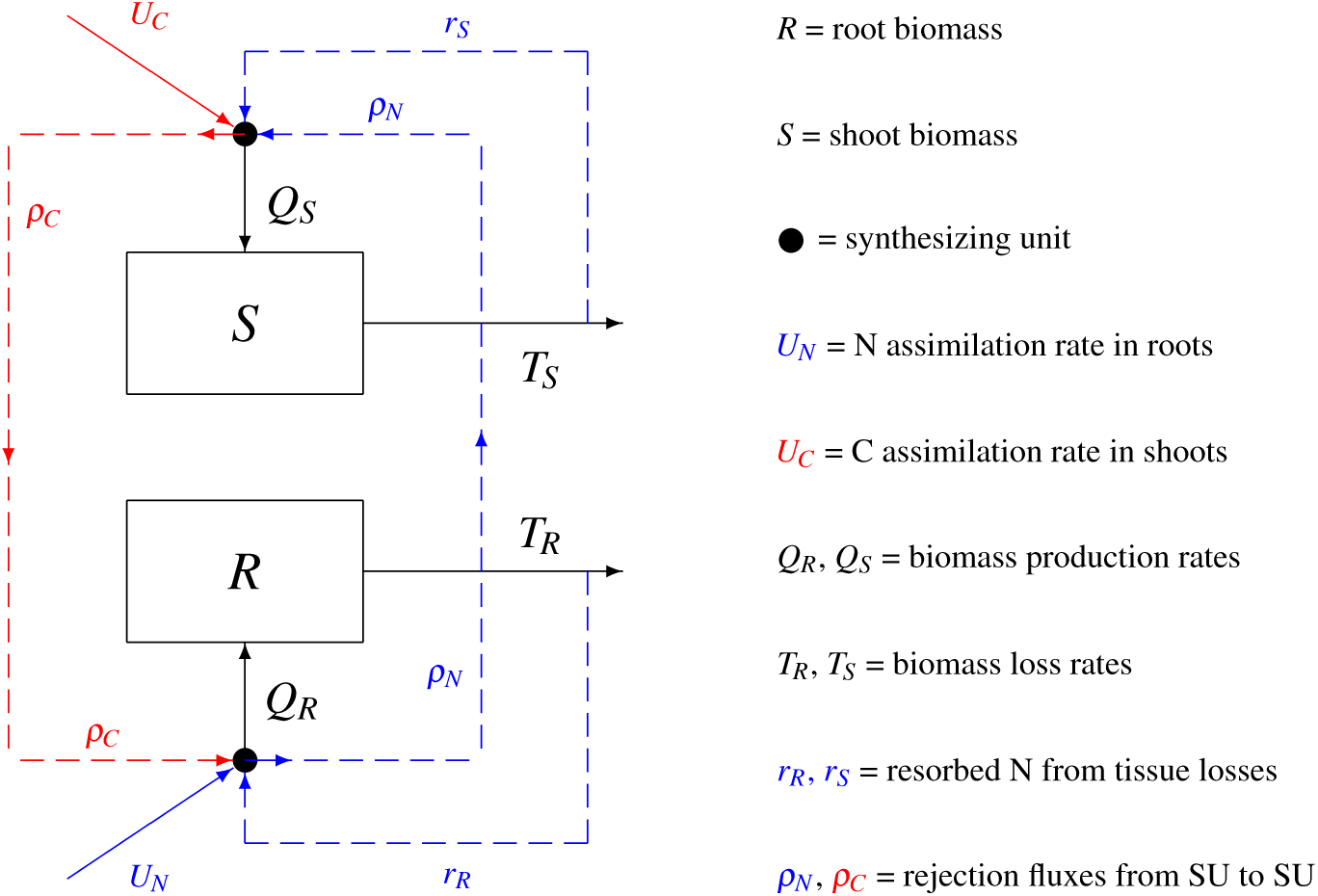
Schematic of model resource flows: C and N arrive at the shoot SU from photosynthetic capture and N rejection from the root SU, respectively; similarly, N and C arrive at the root SU from root assimilation and C rejection from the shoot SU; each SU creates the corresponding plant tissue; shoot and root tissue are lost to herbivory and/or senescence, with some fraction of the N recycled to the local SU.

**Table 1.** List of Symbols (dimensionless unless otherwise specified)

| Symbol | Meaning | Equation |
| --- | --- | --- |
| $F, F_R, F_S$ | Synthesizing unit function (generic, root, shoot) (mass/time) | (14, 15, 16, 17) |
| $h$ | Reciprocal of $k$ in $k$ family of SU functions | (19) |
| $k$ | Exponent parameter in $k$ family of SU functions | (16) |
| $p$ | Scaled relative rate of change of $u$ | (30) |
| $Q_R, Q_S$ | Production rates for roots, shoots (mass/time) | (2, 3, 27) |
| $r_R, r_S$ | Recycling fluxes of roots and shoots (mass/time) | (7) |
| $R$ | Active biomass of roots (mass) | (2) |
| $S$ | Active biomass of shoots (mass) | (2) |
| $T_R, T_S$ | Turnover rates for root and shoot mass (mass/time) | (2, 6) |
| $U_C, U_N$ | Assimilation rates for C and N (mass/time) | (4) |
| $u, u^*$ | Assimilation ratio ($\alpha_C S$)/($\hat{\alpha}_N R$) and equilibrium value | (12, 25, 30) |
| $\hat{u}$ | Modified assimilation ratio ($\hat{\alpha}_C S$)/($\hat{\alpha}_N R$) (includes resorbed N in shoots) | (21) |
| $V(z), W(z)$ | Auxiliary functions for uniqueness | (31, 34) |
| $x, y$ | Auxiliary variables for the symmetric ratio-based SU | (28) |
| $\alpha_C, \alpha_N$ | Assimilation rate constants for C and N (1/time) | (4) |
| $\hat{\alpha}_N$ | Acquisition rate constant for N (1/time) (includes resorption in root) | (8) |
| $\alpha$ | Acquisition rate constant ratio ($\alpha_C/\hat{\alpha}_N$) | (26) |
| $\beta$ | Stoichiometric ratio ($\eta_S/\eta_R$) | (1) |
| $\beta_c$ | Critical value of $\beta$ for guaranteed stability | (32) |
| $\gamma$ | Dimensionless loss rate constant difference $((\gamma_S - \gamma_R)/\hat{\alpha}_N)$ | (26) |
| $\gamma_R, \gamma_S$ | Root and shoot tissue loss rate parameters (1/time) | (6) |
| $\Gamma$ | Contribution of N resorption to shoot SU | (8) |
| $\eta_R, \eta_S$ | N:C ratios for root and shoot formation | (1, 5) |
| $\rho_C, \rho_N$ | Rejection fluxes of C and N (mass/time) | (5) |
| $\sigma_R, \sigma_S$ | N resorption factors for root and shoot | (7) |
| $\Phi(z), \Theta(z)$ | Ratio-based synthesizing unit function and complement | (17, 18, 19, 27, 29) |

1. Root biomass and shoot biomass have fixed, but different, stoichiometries. We define one unit of *R* to be the amount of that component that is made using one mole of C and *η*_*R*_ moles of N; similarly, one unit of *S* is made from one unit of C and *η*_*S*_ units of N. The ratio of the stoichiometric factors,

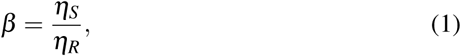

is a dimensionless measure of the relative N:C ratio in shoot formation to that of root formation.
2. Resources are brought into the plant from the environment, with *U*_*C*_ the C-assimilation rate in shoots and *U*_*N*_ the N-uptake rate into roots from soil. These rates are proportional to the shoot and root biomasses, respectively, with rate constants *α*_*C*_ and *α*_*N*_. For convenience of language, we use the term “assimilation rate” for both of these.
3. Production of root and shoot biomasses occurs at “synthesizing units” (SUs), which are idealized production centers that create root and shoot biomass from inputs of C and N. Formulae for production rates at SUs are presented in a separate subsection.
4. Any C input flux to the shoot SU that is not used to produce shoot biomass constitutes a “rejection” flux, denoted by *ρ*_*C*_, that is translocated to the root SU. Similarly, the flux, *ρ*_*N*_, of unused N from the root SU is rejected to the shoot SU.
5. Rejected C from root SU and rejected N from shoot SU are “wasted” resources that are lost to the environment.
6. Root and shoot biomass are turned over at rates (*T*_*R*_ and *T*_*S*_) that are proportional to the relevant biomass, with rate constants *γ*_*R*_ and *γ*_*S*_. Turnover combines a variety of mechanisms, including maintenance (which primarily removes C as CO2), herbivory, senescence, and wood formation.

Fractions *σ*_*R*_ and *σ*_*S*_ of the N used in the original production of lost root and shoot biomass are resorbed and sent to the local SU as an additional input stream. Smaller resorption rates for C can be accommodated in the model by decreasing the net loss rate coefficients *γ*_*i*_, while the larger resorption rates for N are included explicitly. The fractions *σ*_*i*_ can be adjusted to account for variation among species in the fraction of biomass loss that is owing to senescence and in the efficiency of the plant in resorbing N from senesced tissue.

The model dynamics are then given by two biomass balance equations

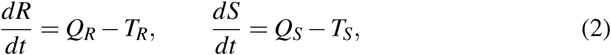

together with a set of equations implementing the model assumptions to yield formulae for the flows in Figure 1:

Biomass production rates:

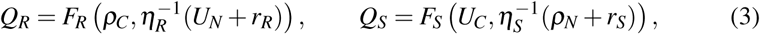

C and N assimilation rates:

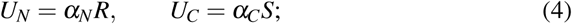

C and N rejection rates:

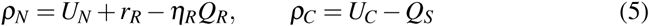

Turnover rates:

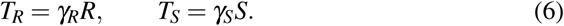

N-resorption rates:

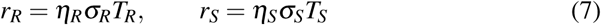

In the production rate equations (3), the functions *F*_*i*_ (*i* = *R,S*) relate the output from the root and shoot SUs to the C and N input rates for component *i*. The stoichiometric factor 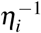 is incorporated into the argument of the function so that the SU models can assume that the correct stoichiometric ratio of the inputs is always 1:1.

We define new compound parameters

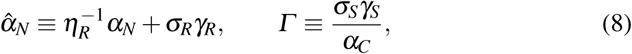

where 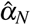 simplifies the notation by incorporating the assimilation and recycling of N in the roots in a single term, while being dimensionally equivalent with the corresponding parameter *α*_*C*_ for the C input to shoots, and *Γ* is a dimensionless measure of the contribution of N recycling in the shoot to shoot SU dynamics. Substituting from (4–7) into (2) and (3) and simplifying with (8) yields the dynamic equations

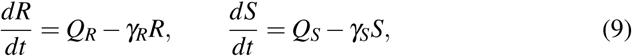

with the production rates determined by the coupled algebraic SU system

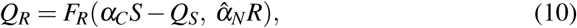

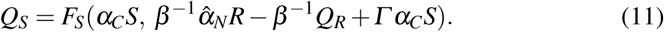

### 2.1 The assimilation ratio

So far we have used the root and shoot biomasses *R* and *S* to represent the state of the system. Because there is no principle of diminishing returns in our model plant, it is a reasonable hypothesis that the system might eventually reach an equilibrium shoot:root ratio *S/R*, with shoot and root growing at a common rate. This suggests modeling the system dynamics using just one of the components as a measure of plant size while using the shoot:root ratio to represent the balance between the organs. We can go one step further by recognizing that the system behavior depends on the assimilation rates achieved by the roots and shoots rather than the biomasses per se. Hence, The best choice of variable to represent shoot:root balance is the “assimilation ratio,” defined as the dimensionless ratio of the C assimilation rate in the shoot to the total N acquisition rate in the root SU (including N resorbed from root turnover as well as N assimilated from the environment):^1^

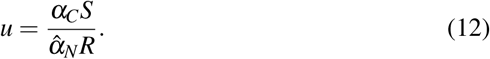

The differential equation for *u* follows from the root and shoot differential equations (2):

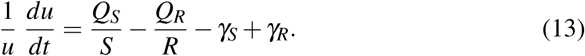

### 2.2 SU Functions

We consider three examples of SU functions, each motivated by different considerations. We first use an idealized SU, the Liebig minimum rule, which assumes both input streams are utilized for biosynthesis to the maximum extent consistent with stoichiometry. This model is obviously an extreme caricature of any plausible biochemical network, since chemical transformations always involve some degree of inefficiency. Nonetheless, we consider it because it allowed us to analytically derive many important qualitative properties of the model that carry over to more realistic representations. Second, we look at an abstraction of the dynamics of a complex biochemical network involved in biosynthesis—the parallel complementary synthesizing unit (PCSU). This was proposed by Kooijman (1998) as a generalization of Michaelis-Menten (MM) enzyme kinetics for situations with two input streams of substrate. We include it because it is a popular representation in studies based on Kooijman’s DEB theory. Finally, we propose a broad class of SU functions that we call the ‘k family of SUs.’. The formula for this family contains a parameter *k* that allows the model to assume any degree of tissue construction “efficiency” (see Subsection 2.2.3), with the perfectly efficient minimum rule as a limiting case.

#### 2.2.1 The minimum rule SU

In mathematical terms, the Liebig minimum rule is represented by the function

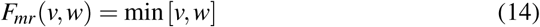

for any two scaled input streams *v* and *w* of C and N, respectively.

#### 2.2.2 The parallel complementary SU

We outline the rationale for the PCSU in Appendix A (online resource), where we show that with one additional simplifying assumption to those of Kooijman, the formula for production rate from a PCSU with input fluxes *v* and *w* is

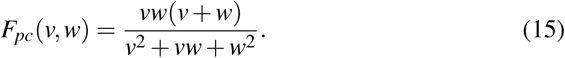

#### 2.2.3 The kSU

We define the “efficiency” of an SU as the biomass production rate when both inputs arrive at unit rate, given that we have defined SUs so that one unit of production requires one unit of each input. The minimum rule has efficiency *E*_*mr*_ = *F*_*mr*_(1, 1) = 1; this is the largest possible efficiency, as the rejection fluxes are both 0. In contrast, the efficiency for the PCSU is noticeably lower, at *E*_*pc*_ = *F*_*pc*_(1, 1) = 2*/*3. In order to explore the ramifications of SU efficiency, we consider an empirically-based family of SU functions (henceforward referred to as the “kSU”) defined by

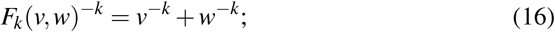

the efficiency of these functions is given in terms of *k* by

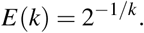

The *k* family can therefore achieve any efficiency from 0 t o 1. The minimum rule SU can be thought of as the limiting kSU as *k* → ∞. The PCSU is not obtainable in the kSU family; however, the kSU offers a very close approximation to the PCSU when the parameter *k* is chosen to match the efficiency *E* = 2*/*3 of the PCSU; that is, *k* = ln 2*/* ln 1.5 ≈ 1.71.

#### 2.2.4 Symmetric ratio-based SU functions

All three of the SU functions we are considering are examples of a broad class of possible SU functions that are symmetric and nonlinear only with regard to the ratio of inputs; that is, they can be written in terms of a function *Φ* such that

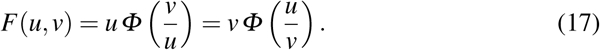

In particular,

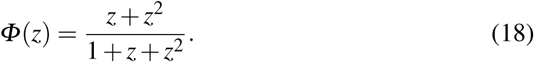

for the PCSU and

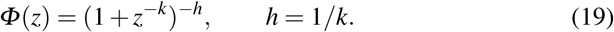

for the kSU.

Given a fixed amount of the resource *u*, any continuously differentiable SU function should satisfy *F*(*u*, 0) = 0, *∂F/∂v* ≥ 0, and lim_*v*_ → _∞_ *F*(*u, v*) = *u*; we therefore assume corresponding properties for *Φ*:

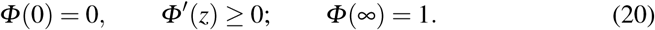

The observation that SUs can be represented by a function of the input ratio alone facilitates the analysis of the model.

### 2.3 Analytical and numerical methods

The model dynamics are not specified solely by the combination of the balance equations and the flux specifications. This is because the production and rejection fluxes are defined *implicitly* with an algebraic system that may have multiple solutions. The analysis necessarily focuses on identifying conditions whereby multiple solutions can occur and elucidating the dynamics of the state variables under these conditions.

Recognizing this inherent difficulty, we adopt two contrasting approaches that allow unambiguous integration of the ODEs even with multiple possible solutions to the algebraic equations, and have verified that, with one caveat mentioned below, both give identical solutions for a broad range of parameter values and initial conditions.

The simplest approach recognizes that it is possible to specify a variant of Euler integration for the differential equations that avoids solving any algebraic equations. Mass balance is achieved by assuming that “transfer” of material between components of the system takes one infinitesimal time step, *dt*. Thus, for example, carbon rejected from the shoot SU at time *t* arrives at the root SU at time *t* + *dt*. Approximating this infinitesimal time step with a very small integration time step *Δt* allows us to update the two state variables, *R* and *S*, using a set of difference equations. Very occasionally this method exhibits numerical instability, but this can be avoided using modified Euler integration.

An alternative approach to assuring a unique solution is to impose an additional problem requirement that the rejection fluxes should be continuous in time whenever possible. This extra condition will be shown to resolve any uniqueness issues that arise in the dynamics. In the case of the minimum rule SU, the algebraic system is simple enough to be solved analytically, allowing a complete description of the dynamics without any numerical analysis (Section 3). In the case of the general continuously-differentiable symmetric ratio-dependent SU, the algebraic SU system and the associated dynamical system can be solved unambiguously over consecutive intervals in time in which the assimilation ratio *u* is monotone (Section 4).

## 3 Analysis and Results for the Minimum Rule SU

We use the term “minimum rule SU problem” to refer to the algebraic problem of obtaining the fluxes *Q*_*R*_ and *Q*_*S*_ from equations (10, 11, 14). The minimum rule SU problem is nontrivial because there may be multiple, positive solutions of these equations. The model dynamics for the minimum rule SU are defined by the differential equations (9) along with the solutions of the minimum rule problem.

### 3.1 Multiple solutions of the minimum rule SU problem

Plant growth with the minimum rule SU depends on whether the plant is C-limited, N-limited, or co-limited by C and N. The classification is based on the current value of a modified assimilation ratio *û*, defined by

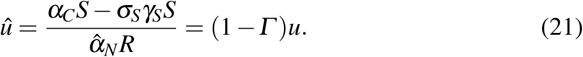

This quantity differs from *u* in that the numerator is the excess rate of C assimilation beyond what is needed to use all of the resorbed N in tissue construction rather than the total rate of C assimilation.

The SU solution has two cases (see online resource, Appendix B, for details), depending on whether *β* is larger or smaller than 1. The two cases coincide at the bifurcation point *β* = 1.

1. If *β* > 1, the C:N ratio for roots is higher than that of shoots, meaning that each resource is relatively more important to the partner that must import it than it is to the partner that assimilates it; in this case the minimum rule SU always has a unique solution, with *Q*_*R*_ and *Q*_*S*_ depending on which resource is limiting.
  a. Both the shoots and roots are C-limited if *û* ≤ *β*^*-*1^ < 1; then all resources are used by the shoot SU:

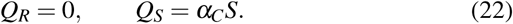
  b. Each component is limited by its imported resource if *β*^*-*1^ < *û* ≤1; then resources are shared:

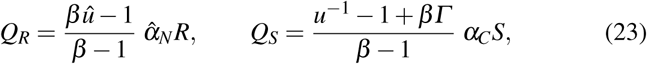
  c. Both the shoots and roots are N-limited if *β*^*-*1^ < 1≤ *û*; then the shoot SU retains its resorbed N and a stoichiometric amount of C, while all other resources are used by the root SU:

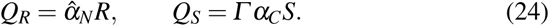
2. If *β* < 1, the relative importance of the local resource for each component is greater than that of the imported resource; in this case the minimum rule SU has multiple solutions whenever *û* is in a range in which the shoots are C-limited while the roots are N-limited.
  a. Both the roots and shoots are C-limited if *û* ≤ 1 ≤ *β*^*-*1^, so all resources are used by the root SU (22).
  b. Each component is limited by its local resource if 1 < *û* < *β*^*-*1^; here the SU system has three solutions (22, 23, 24) and additional rules (see Subsection 3.2) must be used to determine how resources are distributed.
  c. Both the roots and shoots are N-limited if 1 ≤ *β*^*-*1^ ≤ *û*, so the shoot SU uses its own resorbed N and a corresponding amount of C while remaining reources are used by the root SU (23).

### 3.2 Dynamics with the Minimum Rule SU

Growth dynamics are strongly influenced by the possibility of multiple SU solutions; hence, we consider *β* > 1 and *β* < 1 separately.

1. When *β* > 1, the system evolves from any initial state to an equilibrium assimilation ratio *u*\*that is the positive solution of the equation

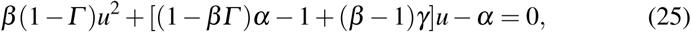

where

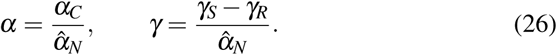
2. When *β* < 1, the dynamics are complicated by the nonuniqueness of SU solutions in the modified assimilation ratio range 1 < *û* < *β*^*-*1^. As noted above, we enforce uniqueness with the additional biologically-motivated requirement that the production rates *Q*_*R*_ and *Q*_*S*_ should be continuous whenever possible, in which case the system evolves to a limit cycle consisting of a phase with *Q*_*R*_ = 0 and *û* increasing from 1 to *β*^*-*1^ alternating with a phase with *Q*_*S*_ = 0 and *û* decreasing from *β*^*-*1^ to 1.

Figure 2 shows examples of the SU solutions and the behavior of the dynamical system for the cases *β* > 1 (top) and *β* < 1 (bottom). Panels (a) and (d) show the extent of root production relative to the amount that would be produced if all N in the root were utilized for root production, while the remaining panels show the dynamics of the assimilation ratio, root biomass, and shoot biomass. When *β* > 1, the assimilation ratio evolves monotonically to its unique stable equilibrium value (25) at a location on the slanted line that is determined by the combination of parameters. There is no root growth when *û* is small and maximum root growth when *û* is large. In the intermediate range, each SU has a surplus of its local resource and must reject some to the other SU. Root growth is near maximum when the C surplus in the shoot is high and near 0 when the C surplus in the root is low.

**Fig. 2.**
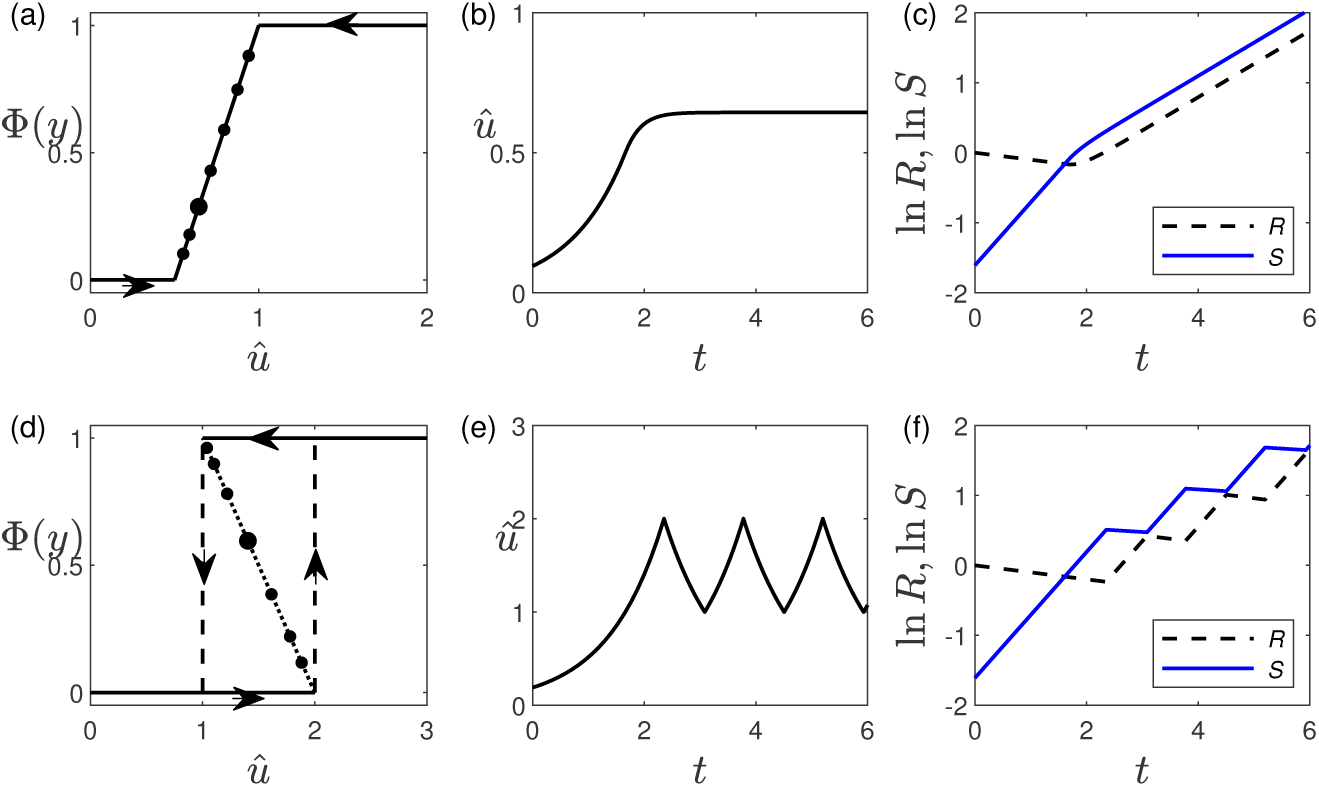
Numerical examples of the two dynamical patterns with the minimum rule SU. Panels (a) and (d): The relative extent of root growth 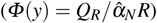 as a function of the modified assimilation ratio *û* (defined in (21)), with *Γ* = 0.05; *β* = 2 in panel (a); *β* = 0.5 in panel (b); and equilibria marked for *α* = 1*/*8, 1*/*4, 1*/*2, 1, 2, 4, 8 (bottom to top). Dotted lines indicate SU solutions that are not realized in dynamic simulations and dashed lines indicate instantaneous changes in the growth rates when *û* reaches its bifurcation values. Panels (b), (c), (e), and (f) show simulation runs with parameters corresponding to the large dots in panels (a) and (d) and with *γ*_*R*_ = *γ*_*S*_ = 0.1.

When *β* < 1, there is never a point at which both SUs are rejecting a surplus to the other. As an example, suppose the initial state is deficient in shoots; that is, *û* < 1, as illustrated in panels (e)–(f). Both components are initially C-limited, so all of the C is used in the shoot SU (*Φ* (*y*) = 0), which causes *û* to increase. The other two SU solutions arise at the moment that *û* = 1; however, both of these solutions (23, 24) produce the result

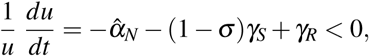

which would move the system back toward lower *û* and the unique solution *Q*_*R*_ = 0, immediately eliminating these solutions. Instead, we must assume that the shoot SU maintains control of the C stream as long as possible; that is, until *û* = *β*^*-*1^. At the moment *û* reaches this critical value, (23) becomes the only viable solution. Shoot production is no longer possible, so the entire C input stream is rejected to the root. Since the system is now N-limited, root production jumps to the maximum. This causes *û* to decrease, but maximum root production stops only when *û* reaches 1, at which point the system state reverses again.

## 4 Analysis and Results for the Continuously-Differentiable Symmetric Ratio-Based SU

The defining equations for the continuously-differentiable symmetric ratio-based SU (10,11) can be recast as

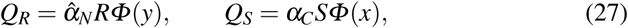

where the auxiliary variables *x* and *y* are determined by a system of two algebraic equations,

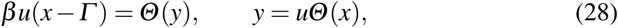

with *u* the current state of the system and

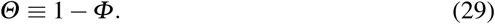

The assimilation ratio dynamic equation (13) becomes

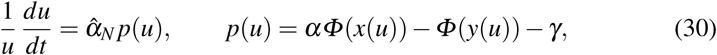

where we have explicitly identified *Φ* in terms of the state variable *u*.

The mathematical analysis is necessarily more complicated than that for the minimum rule SU. Details of the derivations of (27), (28), and (30) and the analytical results summarized in the remainder of this section are in Appendix C (online resource).

### 4.1 Analysis of the continuously-differentiable symmetric ratio-based SU problem

The SU system (28) has these properties:

1. There is at least one solution for any state *u* and parameters *β* and *Γ*.
2. Solutions are unique whenever the SU function satisfies

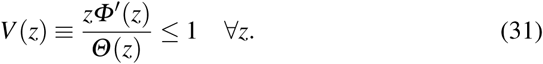
3. When (31) is not satisfied, multiple solutions are possible only if *β* is less than a critical value *β*_*c*_, defined by

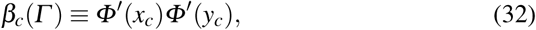

with (*x*_*c*_, *y*_*c*_) defined as the solution of the equations

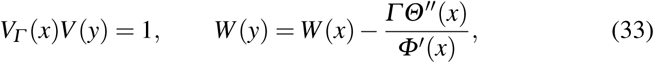

where

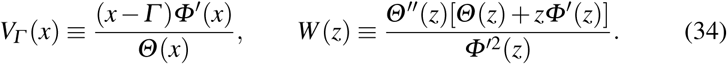

In general, the value of *β*_*c*_ must be determined numerically; it can also be approximated asymptotically for small *Γ* (online resource, (39)).

### 4.2 Growth dynamics with the continuously-differentiable symmetric ratio-based SU

Because production rates are linear functions of component biomass, the model predicts unlimited growth for any viable plant; however, this unlimited growth may occur in such a way that the shoot:root ratio *S/R* and, equivalently, the assimilation ratio *u*, approach equilibrium values. Moreover, the differential equation for *u* is decoupled from the equations for the state variables. These features make the dynamics of the assimilation ratio important to understand. Equilibrium assimilation ratios must satisfy

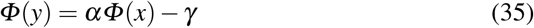

along with the SU equations (28). The growth model has these properties:

1. There is a unique equilibrium assimilation ratio 0 ≤ *u*\*≤ ∞.
2. The equilibrium assimilation ratio is asymptotically stable whenever

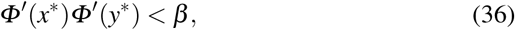

where (*x*\*, *y*\*) is the solution of (35) and

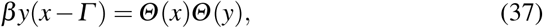

which is obtained by eliminating *u* from (28). This is always true when the SU satisfies (31) or when *β* > *β*_*c*_. When neither of these conditions are met, stability depends on *α* and *γ* as well as *β* and *Γ* (See Figure 3, panels (e) and (f)). **Fig. 3.**
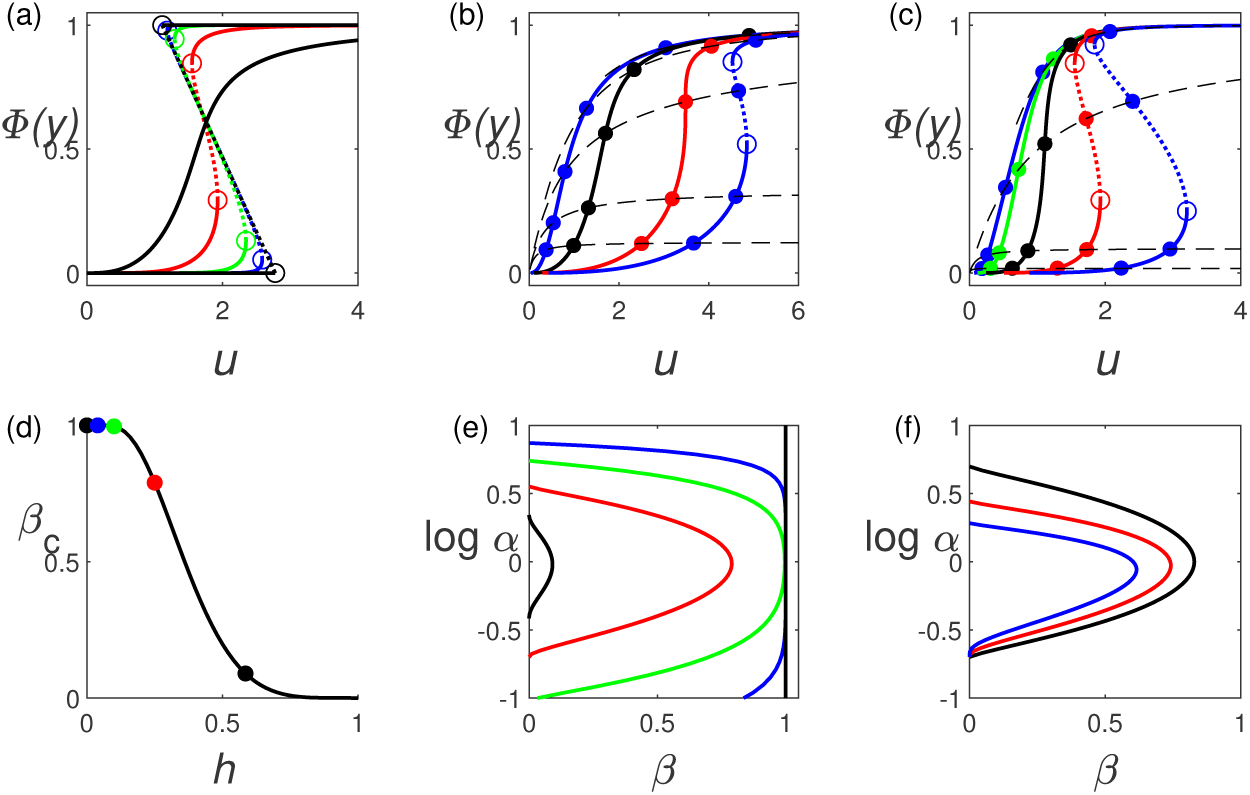
Illustration of the properties of the PCSU and kSU systems. Panels (a)–(c): Solid/dotted curves are the relative extent of root biomass production *Φ* (*y*) as a function of the assimilation ratio *u*, all with Γ = 0.1 (dotted where a potential equilibrium assimilation ratio is unstable); dashed curves are the equilibrium relation (35), all with *γ* = 0; panel (a) is the kSU with *k* = 1.71, 4, 10, 25, ∞ (least to most sigmoidal) and *β* = 0.4; panel (b) is the PCSU with *β* = 2, 0.5, 0.091, 0.05 (left to right) and *α* = 1*/*8, 1*/*3, 1, 3, 8 (bottom to top); panel (c) is the *k* = 4 kSU with *β* = 5, 2.5, 1, 0.4, 0.2 (left to right) and *α* = 0.02, 0.1, 1, 6, 16 (bottom to top). Panel (d): The critical value *β*_*c*_ for the kSU, with Γ = 0.1. Panels (e)–(f): Stability boundaries for the equilibrium assimilation ratio, all with *γ* = 0; panel (e) shows the PCSU, the kSU with *k* = 4, 10, 25, and the minimum rule (left to right), all with Γ = 0.1; panel (f) shows the PCSU with Γ = 0, 0.2, 0.4 (outer to inner).

### 4.3 Results for the PCSU and the kSU

For the PCSU, the inequality (31) is only satisfied for *z* ≤ 1, so uniqueness of solutions depends on having *β* ≥ *β*_*c*_, which is given asymptotically (online resource (39)), with *y*_0_ = 1, *Φ*_1_ = *-*1*/*3, and *Φ*_2_ = *-*4*/*3, as

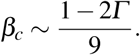

For the kSU, uniqueness is guaranteed when *k*≤ 1 and requires *β* ≥ *β*_*c*_ otherwise; the relationship of *β*_*c*_ and *k* (*h* = 1*/k*) is shown in Figure 3, panel (d).

Additional properties of the SU system are illustrated in Figure 3, panels (a)–(c), as plots of root production relative to its possible maximum 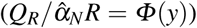 as a function of the current assimilation ratio *u*. Panel (a) shows the effect of SU efficiency on the uniqueness of solutions when *β* = 0.4. The first curve has efficiency of 2/3, matching the PCSU, and shows unique solutions for all *u*. The last curve is the minimum rule SU, showing nonuniqueness over a large range of *u*, and the intermediate curves show the progression from always unique to a large range of *u* with multiple solutions. Panel (b) shows how nonuniqueness develops with decreasing *β* for the PCSU. The third curve from the left in panel (b) has *β* = *β*_*c*_, which is about 0.091 for *Γ* = 0.1; this curve shows a vertical tangent at one point. Any curve with smaller *β*, such as the rightmost curve in the panel, shows an interval of *u* values for which the SU equations have three solutions. More importantly, there is an interval *α*_1_ < *α* < *α*_2_ where the stability criterion (36) is not satisfied; hence, the unique equilibrium assimilation ratio, which is determined by the intersection of the SU curve (28) with the (dashed) equilibrium curve (35) is unstable for moderate values of *α*, where the intersection point is in the dotted portion of the SU curve. Panel (c) is similar to panel (b), but for the kSU with *k* = 4. There are two curves in panel (a) that are the same or nearly the same as one curve in one of the other panels. The least sigmoidal curve in panel (a) is almost identical to the second curve from the left in panel (b), as the behavior of the kSU with *k* = 1.71 is very similar to that of the PCSU. The second least sigmoidal curve in panel (a) is the same as the second curve from the right in panel (c).

The assimilation ratio stability properties of the PCSU and kSU are illustrated in panels (d)–(f) of Figure 3. Panel (d) shows the critical value of *β* corresponding to guaranteed stability, as a function of the parameter *h* = 1*/k*. *h* = 0 is the minimum rule SU, with an efficiency of 0 and *β*_*C*_ = 1. As *h* increases to 1, the efficiency of the SU decreases and the corresponding critical *β* decreases as well. Panel (e) shows the stability bifurcation diagram for the PCSU (leftmost curve), the minimum rule SU (rightmost curve), and the intermediate values *k* = 4, 10, 25; all of these correspond to the large dots in panel (d). Panel (f) shows the effect that *Γ* has on stability. The outermost curve, for *Γ* = 0 is symmetric about log *α* = 1. As *Γ* increases, the unstable region decreases. The decrease is more prominent for large *α* than small because the effect of N resorption is more important when N assimilation is slow. Note that the middle curve corresponds to the *k* = 4 curve of panel (e).

Figure 4 shows the dynamics of the assimilation ratio. The dependence of the equilibrium value on *α* is shown in panel (a). The remaining panels show plots in the *uP* phase plane for the five points marked as solid dots in panel (a). When *β* > *β*_*c*_ (the two lowest curves in panel (a)), *u*\*is a strictly increasing function of *α, P*(*u*) is strictly decreasing, and the assimilation ratio changes monotonically from any initial value to the stable equilibrium. When *β* < *β*_*c*_, there are five possibilities, depending on how the values of *α* and *u*\*compare to the local extrema (*α*_1_, *u*_1_) and (*α*_2_, *u*_2_) (with *α*_1_ < *α*_2_) of *u*\*versus *α*. These five possibilities are shown in the remaining panels. Panels (b) and (c) have *α* < *α*_1_. The stable equilibrium is approached monotonically if starting from *u* < *u*_1_. If starting from *u* > *u*_2_, *u* decreases to *u*_1_; at that point, continuous change in *du/dt* is no longer possible, so the solution jumps to the upper branch and then moves monotonically to *u*\*. Panels (d) and (e) have *α* > *α*_2_, with behavior symmetric to that where *α* < *α*_1_. In the case of moderate values of *α* (panel (c)), the unique equilibrium ratio cannot be reached from either direction; instead, the assimilation ratio achieves a limit cycle with a small range of *u* values and a periodic discontinuity in *du/dt*. Note that the additional rule used to select one of the nonunique SU solutions (trying to maintain continuity of *du/dt*) does not apply when the initial assimilation ratio is between *u*_1_ and *u*_2_. In this case, the correct branch of the *p* vs *u* curve is unknown, as it depends on information from before the initial point.

**Fig. 4.**
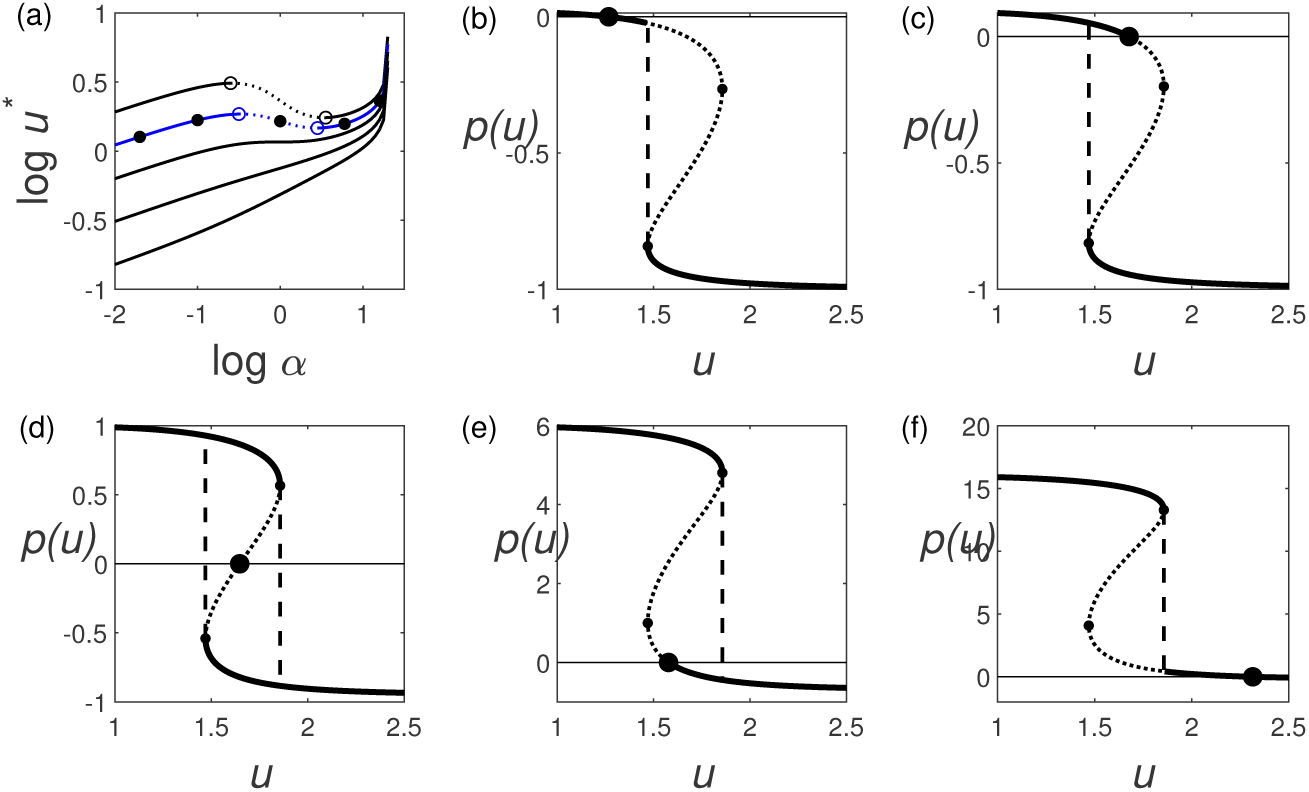
Panel (a): Equilibrium assimilation ratio dependence on the parameter *α* (26) for the kSU with Γ = 0.05, *γ* = 0, and *β* = 0.2, 0.4, 0.811, 2, 5, from top to bottom (*β*_*c*_ = 0.811), with the dotted portions showing equilibrium ratios that are unstable, and dots for the *β* = 0.4 curve for *α* = 0.02, 0.1, 1, 6, 16. Panels (b)–(f): The assimilation ratio increase function *P* (30) with *β* = 0.4, Γ = 0.05, *γ* = 0, and the values of *α* marked with dots in panel (a). The assimilation ratio *u* (12) increases when *P* > 0 and decreases when *P* < 0; hence, trajectories in the *uP* plane are curves that move to the right when *P* > 0, to the left when *P* < 0, and jump to a different part of the curve when continuous movement is not possible.

Figure 5 shows the dynamics of *u* and *R* for the five cases depicted in Figure 4, panels (b)–(f). The different values of *α* are achieved by varying 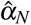; hence, *α* = 0.02 (Figure 4, panel (b)) corresponds to the curves in Figure 5 that show the lowest assimilation ratio *u* and the highest root growth. In panel (c), the bottom curve, for *α* = 0.02, shows a monotone decrease in *u* with a discontinuous derivative, while the second curve, for *α* = 0.1, shows a discontinuous derivative with a change from decreasing *u* to increasing *u*. The top two curves in this panel continuously approach the equilibrium assimilation ratio observed in the corresponding plots of Figure 4. Panel (a) shows corresponding features for the case of a low starting assimilation ratio.

**Fig. 5.**
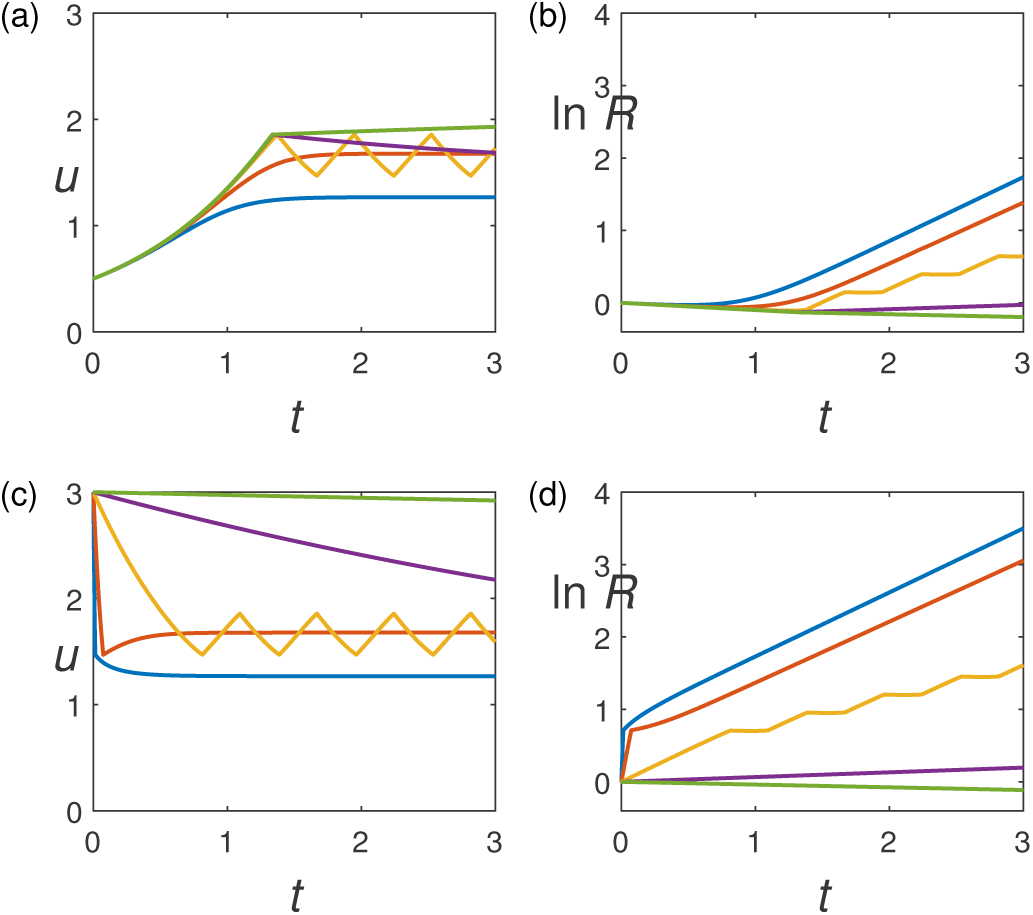
Simulations for the kSU with *k* = 4, *β* = 0.4, Γ = 0.05, *γ*_*R*_ = *γ*_*S*_ = 0.1, *α*_*C*_ = 1, and 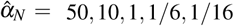, corresponding to the five cases illustrated in Figure 4.

## 5 General Results

Regardless of the choice of SU, the model should be able to predict system behavior that is consistent with general biological principles. Here we consider two issues regarding the response of the system to changes in resource availability or assimilation capacity. Note that we need only consider changes in the dimensionless parameter *α*: either an increase in *α*_*C*_ or a decrease in *α*_*N*_ result in an increase in *α*, and the opposite changes produce a decrease in *α*.

First, we consider the equilibrium shoot:root ratio of the plant changes with *α*; second, we consider the extent to which the local control allocation strategy of the model successfully responds to changes in *α*.

### 5.1 Response of the system to changes in resource availability

Figure 6 illustrates the dependence of the equilibrium shoot-root ratio on changes in the relative availability of C and N. The panels on the left are *S/R*, while those on the right are the ratio of the N contents *SN* = *ν*_*S*_*S* and *RN* = *ν*_*R*_*N*.

**Fig. 6.**
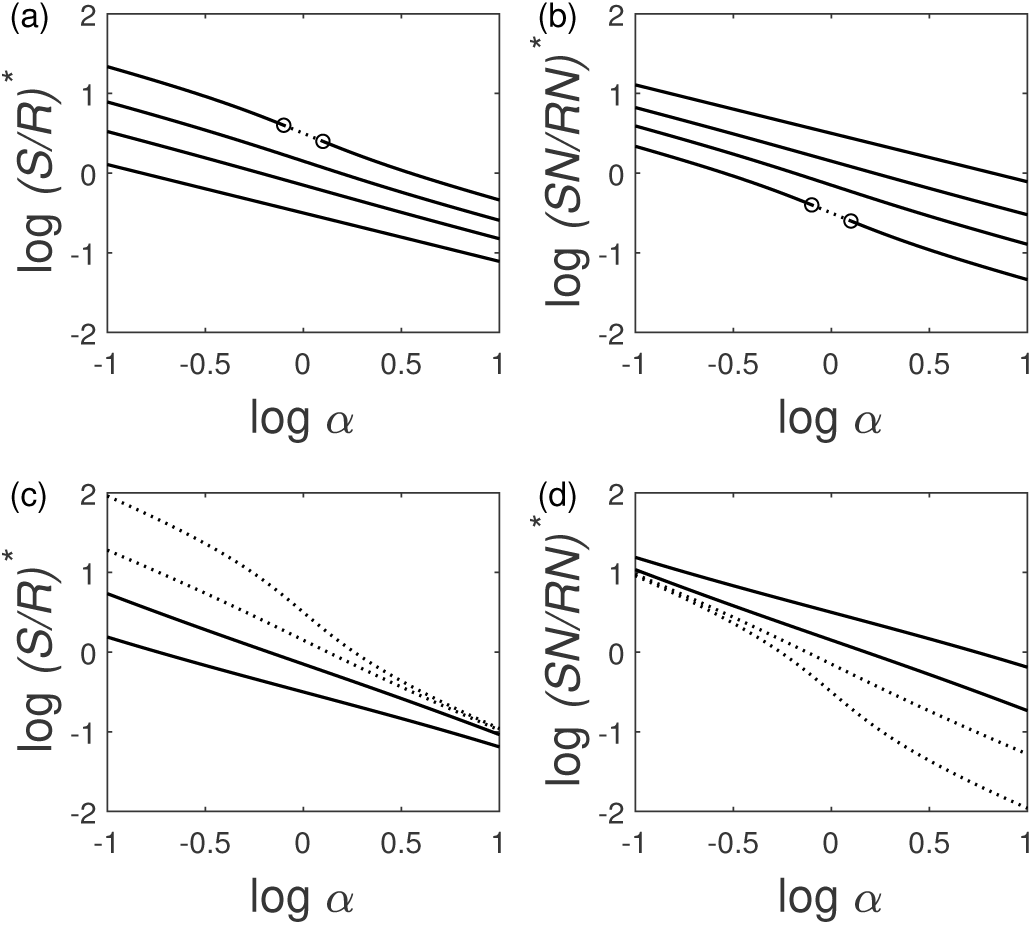
The equilibrium shoot:root ratio for the PCSU (panels (a) and (b)) and the minimum rule SU (panels c and d), with *β* = 10, 2, 0.5, 0.1 (bottom to top on the left panels and top to bottom on the right panels). Panels (a) and (c) show the shoot:root ratio in terms of C content, while panels (b) and (d) use N content.

1. In all cases, decreasing the availability of one of the resources causes a readjustment to build more of the component that collects that resource.
2. A reasonable empirical model for equilibrium shoot:root ratios is

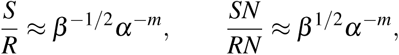

where .6 < *m* < .8 measures the sensitivity of the component balance to variation in resource availability. In general, *m* depends on the SU efficiency and the stoichiometric ratio *β*, and for the unstable range it depends on the assimilation rate constant ratio *α* as well.
3. Larger *β* makes C more concentrated in roots and N more concentrated in shoots, as would be expected from the interpretation of beta as the relative need for the imported resource compared to the local resource. In the specific case *α* = 1, the equilibrium ratios do not depend on SU efficiency.
4. Drivers of instability always increase the response of *S/R* and *SN/RN* to changes in resource availability:
  a. Higher efficiency (higher *k*, leftward on the graph), increases the (negative) slope of the plots.
  b. Lower *β* increases the (negative) slope of the plots.
  c. For the cases where the combination of *k* and *β* permits instability, the curves are noticeably nonlinear, with a larger (negative) slope in the moderate range of alpha.

### 5.2 Optimal balanced growth

The local allocation rule in our model allows a plant to adjust its strategy as resource availability changes, but only in an inflexible way dictated by the behavior of the root and shoot SUs corresponding to the given input streams. It is natural to ask whether the additional flexibility of a global control mechanism might make the plant respond better by controlling those input streams. We can implement global control by introducing allocation parameters *κ*_*C*_ and *κ*_*N*_ to represent the fractions of *C* and *N* that are sent to the local SU. Local control requires that these parameters be unity, but we now postulate the existence of an unspecified global control mechanism that could set *κ*_*C*_ < 1, for example, thereby diverting some of the C resource stream directly to the root. The parameter *κ* is used in a similar way in Cheeseman (1993), except that it is set to a constant value there, while we are allowing it to be chosen dynamically to achieve growth behavior that is optimal according to some specified measure. Total biomass is not necessarily the appropriate measure, as it is unclear whether units of root and shoot should be considered to be of equal value.

While a complete investigation of the optimal control problem is outside the scope of this work, two results suggest the capacity for local control to achieve optimal or near-optimal outcomes.

#### 5.2.1 Optimal balanced growth for the PCSU and kSU

First suppose the function *Φ* satisfies the requirement

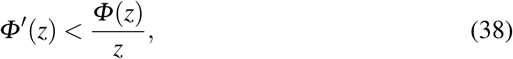

which is the case for both the PCSU and the kSU (see online resource, Appendix D) and that *β* > *β*_*c*_, so that the equilibrium assimilation ratio is stable. When these requirements are met, any pair (*κ*_*C*_, *κ*_*N*_) of fixed allocation parameters results in a stable equilibrium assimilation ratio *u*\*(*κ*_*C*_, *κ*_*N*_), with *u*\*(1, 1) the equilibrium assimilation ratio for local control. The stability of *u*\*means that the root and shoot have a common growth rate *λ* (*κ*_*C*_, *κ*_*N*_), which we take as a working definition of “balanced growth” in the context of our linear growth model. Since this balanced growth rate applies to both root and shoot, there is no ambiguity in defining optimal to be that pair (*κ*_*C*_, *κ*_*N*_) that produces the largest growth rate. We show in Appendix D (online resource) that the largest growth rate under these conditions is always achieved with *κ*_*C*_ = *κ*_*N*_ = 1; hence, the optimal global control strategy for the long term in this case is to use local control.

#### 5.2.2 Optimal approach to balanced growth

It is not surprising that local control tends to achieve the best long-term outcome, as resources diverted directly to the partner are more likely to be wasted than resources sent first to the local SU. The more interesting question is whether local control can be bested by a global control strategy in the approach to balanced growth by making *u* approach the final optimal value *u*\*(1, 1) more quickly than is accomplished with local allocation. Indeed, the theoretical results of Iwasa & Roughgarden (1984) suggest that it would be optimal for the plant to divert resources so as to achieve balanced growth as quickly as possible. However, the question needs investigation because the Iwasa & Roughgarden model is for a system with only one resource.

While a full solution of the optimal control problem is beyond the scope of this paper, it is a relatively simple matter to compare local control against the global strategy suggested above, namely choosing a two-phase approach in which all resources are initially diverted to the deficient partner in phase 1 and then the *κ* values are reset to 1 as soon as the stable equilibrium assimilation ratio *u*\*(1, 1) is achieved. (A strategy consisting of two discontinuous phases is known in control theory as a “bang-bang” strategy – see Hocking (1991) for example.)

Figure 7 illustrates the comparison between a purely local allocation strategy and the two-phase strategy of first shunting all C to the root so as to achieve the ultimate stable equilibrium assimilation ratio as quickly as possible followed by local control to maintain that assimilation ratio. This strategy, shown by the solid curves, is clearly inferior to the purely local control strategy shown by the dashed curves. The short-term benefit of more rapid root growth is not enough to compensate for the inadequate growth of C assimilation capacity otherwise achieved with local control.

**Fig. 7.**
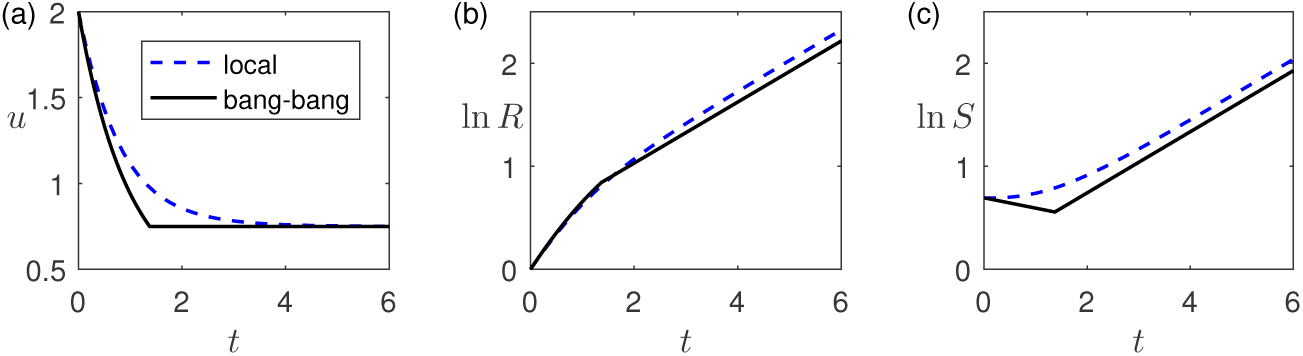
Simulations for the PCSU with 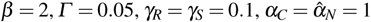; the dashed curves are for local allocation *κ*_*C*_ = 1 and the solid curves are *κ*_*C*_ = 0 until *u* = *u*\*(1, 1) followed by local allocation. The plots show that the overall growth of roots as well as shoots is decreased by a strategy of achieving balance as quickly as possible rather than allowing local allocation to operate.

## 6 Discussion

Allocation of biomass between roots and shoots in plants has often been modeled using some form of global control that optimally allocates resources to maximize the whole-plant growth rate. This approach can be problematic because it assumes fore-knowledge of environmental conditions and the optimal allocation strategy for those conditions, and because the true underlying physiological mechanisms that could achieve such global control, provided it actually exists in nature, are complicated to specify. We show here that such global control is not necessary for mathematical modeling of plant growth and allocation. In the local control theory of plant resource allocation that we present here, each component (shoot or root) is allowed to use as much as it can of its locally produced resource (C or N, respectively), given organ stoichiometry. This mechanism of only sharing surplus resources can be considered a fundamental, higher-level rule of allocation operating between syntrophic entities (i.e., between components within an organism or between organisms). Among plants, it is likely that the lower-level physiological processes achieving this higher-level rule are complex and may vary among plant species, but in our model, these do not need to be specified. This differs from dynamic optimization and global control in that our model does not prescribe what should be maximized, nor does it dictate how a plant should maximize it. Instead, in the local control theory, the optimal outcome emerges from modeling the higher-level rule of only sharing surplus resources, making it more generally applicable. Such purely local allocation rules can achieve the same optimal allocation outcome as a global allocation rule, provided the equilibrium assimilation ratio (the ratio of C assimilation rate to total root N collection rate during balanced growth) is stable, as it is often expected to be in syntrophic systems. Thus, the mechanism of local allocation is sufficient to allow plants to respond to a changing environment so as to maximize growth and allows optimal patterns of root-shoot allocation to emerge from the dynamics of the model, rather than being specified by resource partitioning functions.

### 6.1 Stability and natural systems

Stability of the equilibrium ratio of C assimilation to N assimilation (*u*), and the corresponding equilibrium ratio of shoot biomass to root biomass, depends on three factors: the SU efficiency (1 for the minimum rule, 2/3 for the PCSU, and 2^*-k*^ for the kSU), the relative value of the imported resource to the local resource (*β*), and the resource accessibility balance (*α*), as shown in Figure 3, panels (d)–(f). Below, we describe the conditions under which stability arises in our model, relative to conditions frequently observed in natural systems.

1. Stability is guaranteed when the SU efficiency is very low.
2. Stability is guaranteed when the imported resource is of greater value to each component than is the locally-produced resource (that is, the C:N ratio of the N-producing component is higher than that of the C-producing component).
3. When neither of these two sufficient conditions holds, there is a threshold value of the stoichiometric ratio for shoots and roots (*β*) for stability that increases from 0 for very inefficient SUs to 1 for the maximally efficient SU (the minimum rule SU).
4. Irrespective of SU efficiency, the equilibrium assimilation ratio can still be stable if there is sufficient imbalance in the availability of resources (Figure 3, panels (e)–(f)).

The second of these conditions is perhaps the most important, since it describes precisely the context under which there would be natural selection for symbiosis, and so is often expected to be met in syntrophic systems. In vascular plants, the N requirement for C assimilation is considerably larger than for N uptake, which implies a large value for *β*. The importance of this point is reinforced by noting that nitrogen is not necessarily the limiting resource represented by “N.” For example, carbon fixation can be limited by the regeneration of ribulose bis-phosphate (Mott et al. 1986). But again, this would make the C:N ratio of shoots lower than that of roots (with phosphorus as N). For syntrophies in which the components have lower need for the imported resource, *β* may be less than one, but perhaps not low enough to permit instability.

Should none of the three sufficient conditions for stability be met, the deciding factor is the dissimilarity of assimilation rate coefficients *α*_*C*_ and *α*_*N*_ for photosynthate and inorganic nutrient respectively. These coefficients represent a composite of plant traits, such as specific leaf area, specific root length, and C:N ratio, which determine the assimilation efficiency of a unit of shoot or root biomass (Reich et al. 2003), as well as environmental factors, such as the availability of sunlight and soil nutrients. Large differences in the assimilation rate coefficients push the system toward stability, whereas small differences push it toward instability. Our model treats these coefficients as constants, but in a real system they may vary over time as plants plastically grow organs with different trait values or as environmental conditions change. Thus, a system that is sufficiently efficient and has stoichiometry sufficiently favorable for instability might alternate between periods of stability and instability. Separating plant trait-related factors and environment-related factors into different parameters would allow greater flexibility in modeling the interactions between components of specific syntrophic systems and their plastic responses to environmental variation.

When the equilibrium assimilation ratio *u*\*is unstable, the system oscillates between periods when a large root construction rate causes the assimilation ratio to decrease beyond *u*\*and periods when a large shoot construction rate forces it to increase beyond *u*\*(see Figure 4, panel (d)). The system therefore cycles between conditions corresponding to mutualism (simultaneous sharing of locally produced resource) and parasitism (hoarding of the locally produced resource while still receiving the imported resource), analogous to the concept of the mutualism-parasitism continuum (Johnson et al. 1997; Bronstein 2001). If it has enough of the imported resource, each SU will consume all of its local resource, causing rapid growth while inhibiting its partner’s growth. If this trend is unchecked, the assimilation ratio will move beyond its stable value and the relationship will become parasitic. However, this condition in our syntrophy model is self-correcting. The more the growth of the parasitic partner, the more of the local resource it has available; eventually there is so much that some must be shared due to tissue stoichiometry. Instability occurs when each of the components can acquire resources fast enough, due to their comparable alphas, to take a turn as the parasite. This alternating parasitism is checked, however, when either of the two sufficient conditions for stability (1 and 2 in the list above) are satisfied. While the true biological mechanisms underlying the mutualism-parasitism continuum are of course more complicated than those in our model, the context-dependency of stability in resource-sharing in our model corresponds to what is observed in natural symbioses (e.g., Denison & Kiers 2004).

### 6.2 Local versus global control

Any model of plant growth needs to have rules for allocating resources to the different organs that comprise the plant. It is intuitive to prefer allocation outcomes to be optimal in some sense, such as producing a maximum rate of biomass growth. This is the premise of Iwasa & Roughgarden (1984), as generalized by Velten & Richter (1995), which identified the allocation strategy that achieves optimal growth of a plant model in which carbon and water are the only scarce resources. In general, one might expect the greater flexibility of global control to yield evolutionarily superior outcomes than can be produced via local control. This is not the case in our local control theory of plant resource allocation, as we demonstrated by comparing our standard model with a variant that incorporates global control by allowing time-dependent fractions of locally-produced resources to be sent directly to the partner without having to be rejected by the local SU. In the long-term simulation (with parameters such that the equilibrium assimilation ratio is stable), shunting any fixed portion of either resource directly to the partner, as is assumed for above vs belowground carbon allocation in some in some dynamic global vegetation models (e.g., Ostle et al. 2009), rather than sharing only the rejection flux, always decreases the long-term growth rate, compared to local control. Thus, local control produces optimal growth in the long term.

The balanced growth (or functional equilibrium) hypothesis states that exogenous resource collection rates always match the weighted average stoichiometry across all of the different molecules that comprise the plant (Shipley and Meziane 2002). This hypothesis only applies under the restrictive assumptions of a constant environment and no damage to the plant that would upset the balanced resource collection capacities. However, plants often experience dramatic changes in the ability to acquire resources, either due to changes in the availability of resources or assimilation capacity (e.g., defoliation by a pest). The local control model states that the way to restore the balance is to allocate endogenous resources so as to increase the collection capacity for the deficient exogenous resource, without decreasing the collection rate of the excess resource. Moreover, in the absence of resource storage, intuition suggests that the best short-term strategy in such cases would be to allocate all resources to growing the deficient organ until the optimal ratio of resource assimilation capacities is achieved. This strategy is optimal in the simpler setting of Iwasa & Roughgarden (1984) and Velten & Richter (1995), but it does not perform as well as local control in our model. The initial situation for the simulation presented in Figure 7 is the aftermath of a sudden loss of N assimilation capacity. The two-phase strategy of initially shunting all C directly to the root has the short-term benefit of maximizing immediate root growth. However, the cost is that the plant fails to invest any resources into maintaining its C assimilation capacity. The more rapid loss of this capacity that occurs with the two-phase strategy as compared to local allocation has a long-term detrimental effect on root growth by decreasing the future flow of excess C from shoot to root. Local control seems to find the right balance between these benefits. This result illustrates the concept of the time-value of uptake capacity, in which earlier investments in uptake capacity yield greater uptake in the long-term due to the compounding effect (Lerdau 1992). While this concept has previously been applied to leaves (Westoby et al. 2000), our model suggests that it should apply equally well to roots, as a scenario in which there is a sudden loss of C assimilation is analogous.

The equilibrium assimilation ratio is always optimal; however, it can only be achieved through local control when it is stable, such as when *β* > 1, which should usually be the case in natural systems like most vascular plants. When the ratio is unstable, the maximal growth rate can only be achieved through global control. The optimal behavior in this case appears to be to use local allocation until the equilibrium assimilation ratio is achieved and then use global control to maintain that assimilation ratio. Thus, the success of local control in managing resources in a theoretical plant model in most cases suggests that models of plant growth do not require an allocation submodel that assumes global control and considers strategies to achieve some optimal outcome.

### 6.3 Assumptions, caveats, and extensions

The local control theory of plant resource allocation developed here makes several simplifying assumptions that may need to be relaxed in application to real biological systems. Realistic mortality mechanisms would need to be incorporated in order to provide insights on the adaptive value of different allocation strategies that involve resource storage, defense, stoichiometric plasticity, or dormancy. As is true of many plant growth models, complicated processes have been abstracted into single parameters. For example, the *σ*_*S*_ parameter that controls the fraction of N in leaf turnover that can get recycled is actually a combination of resorption efficiency (Aerts, 1996) and herbivory. The former is a functional trait, while the latter is a combination of ecological conditions and the extent to which resources are allocated to chemical or physical defenses. To fully understand the complex determinants of trade-offs between allocation to growth versus defense, the resorption efficiency and herbivory factors should be decoupled, but the latter would instead need to be coupled to the assimilation coefficient *α*_*C*_ to capture growth-defense trade-offs (Herms and Mattson 1992); i.e., more resources used for defense against herbivory means less resources used for the machinery needed for C assimilation.

The interacting components (roots and shoots) in the local control model presented here are characterized as “biomass” with fixed stoichiometry. Actual plants display considerable plasticity in nutrient concentrations of their organs, in response to changing environmental conditions (Rozendaal et al. 2006). Moreover, both roots and shoots may contain energy or nitrogen-rich storage compounds that require minimal maintenance (Chapin et al. 1990). As a result, precise quantitative model predictions, such as critical values for a model parameter, should not be viewed as absolutes. However, we suggest that qualitative patterns, especially trends in response to changes in environment, represent robust predictions.

The scope of our model is necessarily limited. Consistent with the spirit of many ecological models, we use an idealized abstraction of a real plant as having only two components (root and shoot). We focus exclusively on growth, not on reproduction or survival. We do not explicitly model competition between plants for light or nutrients, although it would be straightforward to develop an individual-based population model in which each plant followed the rules proposed in our model but was coupled to a shared environment.

### 6.4 Implications for modeling syntrophic mutualisms

The local control theory of plant resource allocation that we have developed here can be applied or adapted to a wide variety of obligate syntrophic relationships, including mutualisms among organisms, particularly since local control as a mechanism for resource allocation between partners implicitly assumes that the unit of selection is the partner, not the holobiont. Mycorrhizal fungi are obligate symbionts of plants that are largely incapable of acquiring carbohydrates on their own, but are more efficient than plant roots in acquiring nutrients from soil. They trade soil- and litter-derived nutrients, such as phosphorus, for plant-derived carbohydrates (Smith & Read 2008). Our model would require some modification in order to be applied to plant-mycorrhizal interactions, since it assumes that each partner is incapable of acquiring the resource it imports from its partner. However, mycorrhizal fungi produce structures inside plant roots that absorb carbon and structures outside of the plant root that absorb resources from soil. Allocation to such intra-versus extra-radical structures is plastic for arbuscular mycorrhizae and depends on competitive interactions with other arbuscular mycorrhizae inside the plant root (Engelmoer et al. 2011), which is analogous to the root-shoot system in our model.

There are also many examples of syntrophic symbiosis in microbial systems, in which one bacterial population provides an obligate requirement of another. Yeoh et al. (1968) reports results from interacting bacterial populations, in which each supplies a vitamin requirement for the other, but one also produces an inhibitor. There were large amplitude oscillations in continuous culture of these bacteria, superficially resembling those in Figure 2, panel (e). Adding a proteolytic enzyme (presumably destroying the inhibitor) lead to a growth burst stimulated by the mutualisitic interaction, followed by decline as the enzyme washed out, with oscillations ultimately resuming. We speculate that the dynamical mechanisms causing oscillations in this mutualist/inhibitor system resemble those analyzed in this paper, with Figure 3 panels (a)–(c) showing inhibition of root biomass production in response to shortage of photosynthate. Likewise, Weederman et al. (2013) developed a model of anaerobic digestion with syntrophy and showed that inhibition may introduce regions with multiple steady states and may stabilize some equilibria. A multi-species generalized Lotka-Volterra model with both positive (mutualist) and negative (inhibitory) terms explains observed experimental results defining the effects of antibiotics on *Clostridium difficile* infections in mammals (Jones & Carlson 2018).

The local control theory of plant resource allocation presented here is similar to the Muller et al. (2009) DEB model of corals, with the shoot analogous to the algal symbiont (the source of photosynthate) and the root analogous to the animal host (the source of nitrogen). In that model, the interspecific interaction involves sharing the surplus and, with parameters chosen as representative of a scleractinian coral, the symbiont to host biomass ratio stabilizes, as in our model. However, the Muller et al. (2009) model has other potentially stabilizing processes, including intraspecific processes involving energy reserves for each species. Whether these processes, local allocation, or a combination of these, is responsible for the stability of that model is not clear. However, the analysis in the present paper points clearly to sharing the surplus as the primary stabilizing mechanism. Furthermore, our analyses indicating no need for global control of resource allocation in plants gives added credibility to the conclusion by Muller et al. (2009) that active mechanisms, such as the digestion or expulsion of symbionts, are not necessary for a relatively stable symbiont density. Our model shares other properties with more complex counterparts. For example, Muller et al. (2009) showed that the symbiont to host ratio decreases with increasing irradiation (corresponding to increasing *α*_*C*_) and increases with increasing availability of dissolved inorganic nitrogen (corresponding to increasing *α*_*N*_), consistent with the simpler model here (Figure 6). Cunning et al. (2017) cites many observations of negative trends between irradiance and symbiont density and one study in which increasing symbiont to host ratio increases with increasing dissolved inorganic nitrogen in the environment.

Looking forward, there is a growing literature with applications of models incorporating syntrophic interactions to societally important challenges, such as optimizing plant-microbial interactions in agricultural settings to maximize crop yields and modeling the growth dynamics of infectious agents confronted with antibiotic inhibitors. Environmental change can also impact plant-microbe interactions within a plant (e.g. nitrogen fixation), leading to non-monotone dose-responses to a contaminant (Priester et al, 2012, 2017; Klanjscek et al 2017) caused by the interplay of the demands of energy-intensive N-fixation with the benefits of enhanced supply of nutrient. The work reported in this paper points to a potential role for mechanistic models of syntrophy where the observed outcome (apparent mutualism versus parasitism) emerges from the dynamics and is not assumed from the beginning.

## Appendix A: The Parallel Complementary Synthesizing Unit (PCSU)

In this appendix, we outline the rationale for the parallel complementary SU model, proposed by Kooijman (1998) and widely used in Dynamic Energy Budget (DEB) theory (e.g. Kooijman, 2010). We have added this appendix because descriptions in the literature known to us are couched in the terminology and concepts of DEB theory, which is likely to be unfamiliar to readers of this paper.

We aim to derive a formula for the function *F*(*v, w*), the rate of production of biomass from two input streams of elements *V* and *W*, and denoted by *v* and *w*, with the input rates already scaled so that one unit of biomass is made from one unit of each element. The SU is a caricature representation of the network of reactions involved in biosynthesis as a “generalized enzyme” that can be in one of four states: “free”, bound to one unit of V, bound to one unit of W, and bound to both one unit of V and one unit of W, in proportions denoted by *θ*_••_, *θ*_*v*•_, *θ*_•*w*_ and *θ*_*vw*_, respectively. Figure

**Fig. 1.**
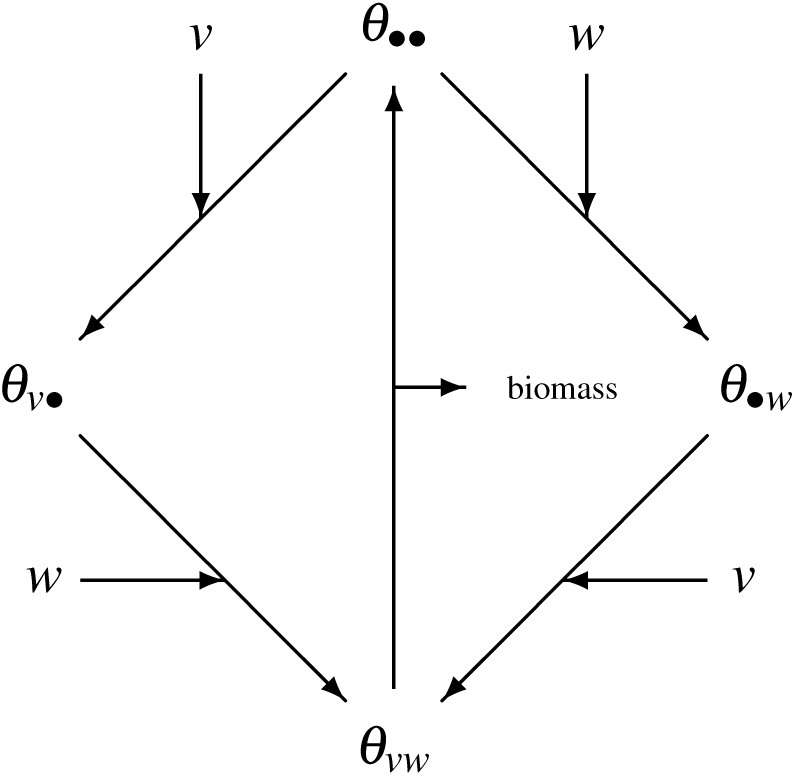
A schematic diagram of the PCSU.

A1 shows the possible transitions among states. Transition rates into states that accept inputs of V or W are assumed proportional to relevant inputs. The transition rate out of state *θ*_*vw*_ is denoted by *H*^*-*1^, where *H* represents the mean time that a SU spends in state *θ*_*uv*_. The dynamic equations are then:

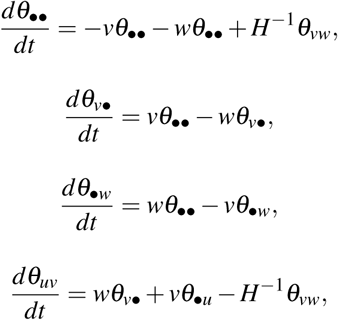

with

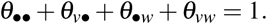

The rate of of biomass production is *F*_*pc*_(*v, w*) = *H*^*-*1^*θ*_*vw*_. If we assume that the time scale of transitions among states is much faster than the time scale of whole organism dynamics (the focus of this paper), then the above dynamical system is in pseudoequilibrium. This permits explict calculation of the function *F*_*pc*_ as

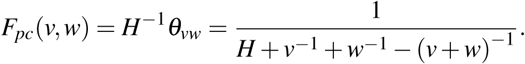

If we further assume that the final transition from *θ*_*vw*_ is much faster than the others, i.e. *H* → 0, then

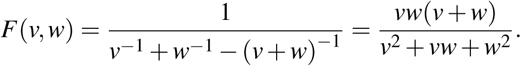

## Appendix B: Analysis of the Minimum Rule SU

### B.1 Existence and uniqueness of SU solutions

The minimum rule SU problem

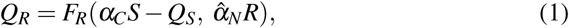

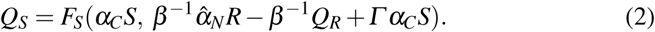

is conveniently recast in terms of the rejection fluxes

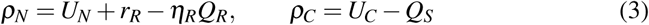

as

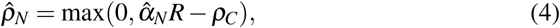

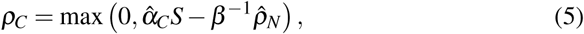

where

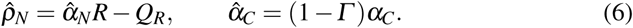

The solution of this problem is best worked out in separate cases.

#### B.1.1 Case 1: Shortage of C (ρ_C_ = 0)

The N rejection flux follows immediately from (4):

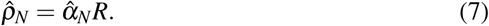

However, this solution is consistent with (5) only if 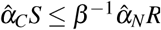, or

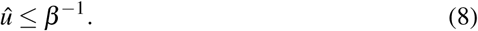

#### B.1.2 Case 2: Shortage of N 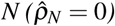

Analogous to Case 1, the C rejection flux (from (5)) is

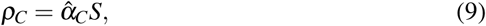

which is consistent with (4) only if

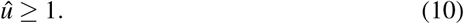

#### B.1.3 Case 3: No local shortages 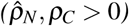

In this case, the SU equations become

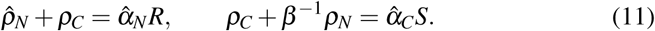

Assuming *β* ≠ 1, the solutions are

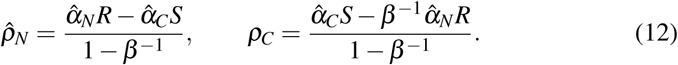

(The special case *β* = 1 requires 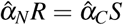 and results in 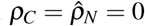. This case has no practical relevance, as the probability that the real number *β* is *exactly* 1 is 0.) This solution is valid only when both rejection fluxes are positive, which corresponds to the interval *β*^*-*1^ ≤ *û* ≤ 1 when *β* > 1 and the interval 1 ≤ *û* ≤ *β*^*-*1^ when *β* < 1.

The results of all three cases are combined to make the full SU solution profile, described in Subsection 3.1 of the main paper.

### B.2 Dynamic behavior

The dynamic behavior of the minimum rule system depends on whether *β* is larger or smaller than 1. We consider these cases separately.

#### B.2.1 *β* > 1

Suppose the system begins with *û* < *β*^*-*1^. This is a low enough assimilation ratio that the SU solution has *Q*_*R*_ = 0 and

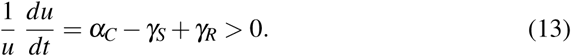

The assimilation ratio increases at this rate until *û* = *β*^*-*1^. Because the quantities *Q*_*R*_ and *Q*_*S*_ are continuous, the derivative *du/dt* is continuous as well; hence, the assimilation ratio increases into the interval *β*^*-*1^ < *û* < 1. In this interval, we have

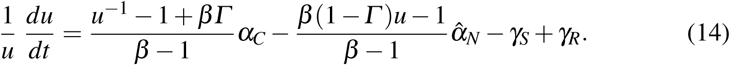

Setting *du/dt* = 0 yields the equilibrium equation

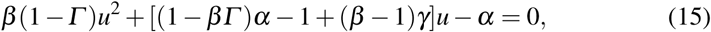

which has a unique solution *u*\*∈ {(1 − *Γ*)^*-*1^*β*^*-*1^, (1 − *Γ*)^*-*1^}. The solution is clearly stable because (1*/u*)*du/dt* is monotone decreasing in *u*. Similarly, an initial assimilation ratio larger than *u*\*results in a monotone decrease from the initial assimilation ratio to the equilibrium. Hence, the equilibrium is globally asymptotically stable.

#### B2.2 *β* < 1

Each of the three cases (1, shortage of C; 3, shortage of neither; 2, shortage of N) has a corresponding differential equation for the assimilation ratio:

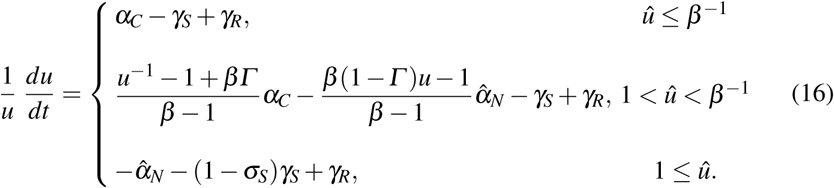

Now suppose the system starts with *û* < 1. The first of the assimilation ratio equations holds, and *û* increases unless *α*_*C*_ is so small that the plant is not viable. Once *û* = 1, all three SU solutions are possible, but only the Case 1 solution yields a continuing increase in *û*. On biological grounds, the resolution of the lack of mathematical uniqueness should be that the system tries to maintain a continuous growth rate, which can only happen if it remains in Case 1 beyond *û* = 1. The system will remain in that case until *û* = *β*^*-*1^, when Case 1 is no longer viable. At this point, there becomes an infinitesimal N shortage, so the system must jump to Case 2. From here, *û* must decrease; hence, all three cases give solutions to the SU equations, but the biological principle of continuity of growth selects Case 2 and *û* decreases until it reaches 1, at which point the cycle begins again.

If the system starts with *û* > *β*^*-*1^, then the behavior is analogous, with Case 2 necessarily holding at the beginning and continuing until *û* = 1, at which point we are in the same limit cycle as above. If the system starts in the intermediate range 1 <, *û* < *β*^*-*1^, then the particular Case must be specified as part of the initial condition. Cases 1 and 2 would be on the stable limit cycle, whereas Case 2 would evolve toward one of the extreme values for *û*. Case 2 has an equilibrium solution with the same formula as in the *β* > 1 case, but with *du/dt* an increasing function of *u* (since the denominators in the formula are now negative), this equilibrium is unstable.

## Appendix C: Analysis of the Continuously-Differentiable Symmetric Ratio-Based SU

For any continuously-differentiable symmetric ratio-based SU

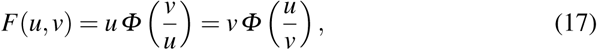

the SU system (1,2) can be recast in terms of a “shoot input ratio” *x* and “root input ratio” *y*, defined by

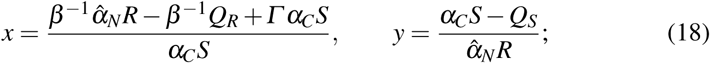

the defining equations (1,2) then become

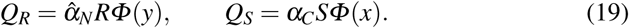

Using these formulas to eliminate *Q*_*R*_ and *Q*_*S*_ from (18) reduces the SU system to a pair of equations for *x* and *y*:

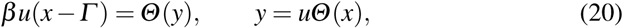

where

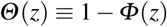

and *u* is the assimilation ratio

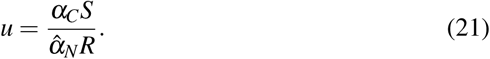

Replacing *Q*_*R*_ and *Q*_*S*_ using (19) changes the dynamic equations for the state variables to

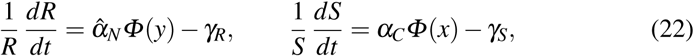

and the equation for the assimilation ratio *u* to

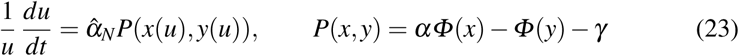

where

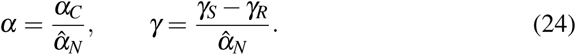

In this formulation, the assimilation ratio equation is decoupled from the root and shoot equations, allowing for study of its long-term behavior in terms of the parameters *α, β, Γ*, and *γ*.

### C.3 Existence and Uniqueness of SU Solutions

#### C3.1 Proof of existence

Given a fixed value of *u*, we can reduce the SU system (20) to a single nonlinear equation

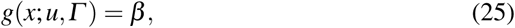

where (suppressing the parameters *u* and *Γ*)

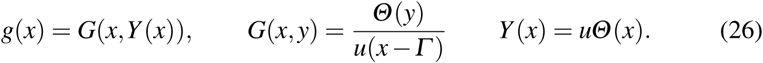

The properties of the SU function *Φ* ensure that *Y* → 0 as *x* → ∞, so that the numerator of *g* goes to 1 while the denominator goes to ∞; hence, *g* → 0 as *x* → ∞. Meanwhile, as *x* → *Γ*, the denominator of *G* goes to 0 and *Θ* (*Y*) > 0, so *g* → ∞. Since *g* is continuous, it therefore achieves all positive values; hence, the equation *g*(*x*) = *β* has at least one solution.

#### C.3.2 Sufficient condition on Φ for uniqueness

With *u* given, the SU solution is guaranteed to be unique if *g*^′^ ≤ 0 for all *x*. If *u* is unspecified, this condition generalizes to

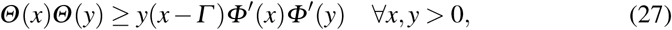

where *x* and *y* are constrained by (20) to satisfy the equation

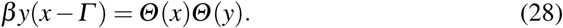

The condition (27) can be rearranged as

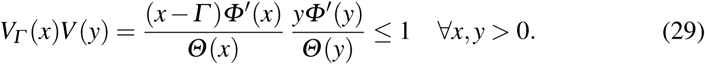

This condition always holds whenever *V* (*z*) ≤ 1 for all *z* > 0.

#### C3.3. Sufficient condition on β for uniqueness

Even if the sufficient condition on *Φ* for uniqueness is not satisfied, multiple solutions to the SU equations are possible only for values of *β* for which (28) has solution pairs (*x, y*) such that

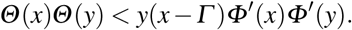

Such values will occur as intervals bounded by solutions of (28) that satisfy

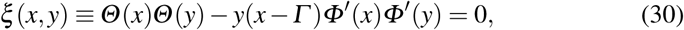

in which case we can replace (28) with the equivalent equation

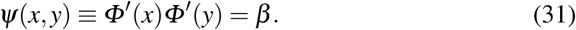

In the limit *y* → 0, the SU properties imply *Θ* (*y*) → 1, in which case (28) implies *Θ* (*x*) → 0. This in turn implies *x* → ∞, whence *Φ*^′^(*x*) 0. Given that *Φ*^′^(*y*) is bounded, we have *ψ* →0 as *y*→ 0. Hence, multiple solutions to the SU equations can only occur on an interval 0 < *β* < *β*_*c*_, where *β*_*c*_ is the maximum value achieved by *ψ* subject to the constraint *ξ*= 0.

The maximum of *ψ* on the curve *ξ*= 0 must occur at a point that satisfies the Lagrange multiplier rule; given that ∇*ψ* ≠ 0, the Lagrange multiplier can be eliminated to yield the equation *ψ*_*x*_*ξ*_*y*_ = *ψ*_*y*_*ξ*_*x*_, or

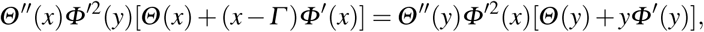

which can be written as

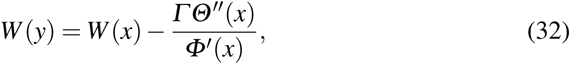

where

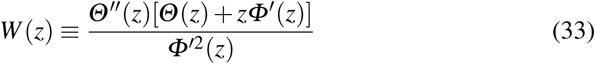

(using first derivatives of *Φ* and second derivatives of *Θ* because these quantities are conveniently nonnegative). In general, *β*_*c*_ can be found numerically from the original optimization problem or by finding the solution (*x*_*c*_, *y*_*c*_) of the system (30, 32) and calculating *β*_*c*_ = *ψ*(*x*_*c*_, *y*_*c*_) from (31).

#### C.3.4 Asymptotic calculation of β_c_

Under the reasonable assumption that *Γ* is small, corresponding to the biological assumption that resorption of N is only a small fraction of N assimilation, we can obtain an asymptotic solution of the form

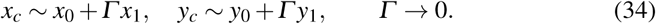

For convenience, we rewrite (30) as

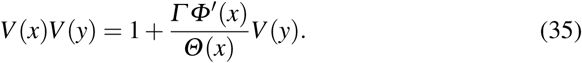

To leading order, the system (32, 30) becomes the symmetric system

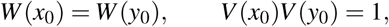

which has the (unique if *W* is monotone) solution *x*_0_ = *y*_0_ with

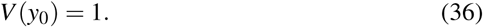

This last result can also be written as

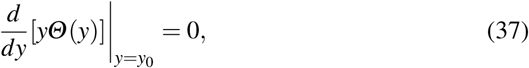

allowing the interpretation of *y*_0_ as the point where the function *yΘ* (*y*) achieves its maximum value.

Defining *Φ*_1_ and *Φ*_2_ by

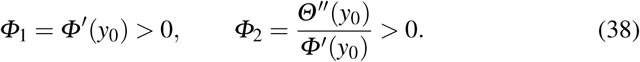

and using the results of the leading order approximation, we obtain the two-term approximations

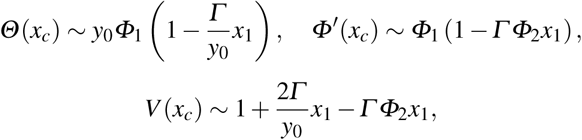

with similar approximations for *y*_*c*_, and

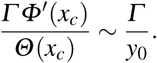

Substituting the last two of these into (35) yields

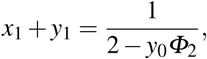

whence

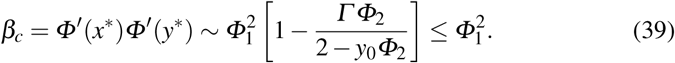

### C.4 Dynamic behavior of the assimilation ratio

#### C4.1 Existence of a unique equilibrium assimilation ratio

Equilibria for the assimilation ratio equation (23) are defined by

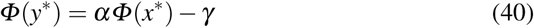

along with the SU equations

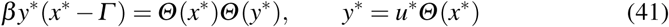

We can think of (41) as defining functions *y*\*(*x*\*) and *u*\*(*x*\*), whence the equilibrium equation (40) then identifies equilibria by defining *x*\*as a function of *α*. As *x* → ∞, we have *y* → 0 and so *P*(*x, y*) → *α* + *γ*. Similarly, as *x*→ *Γ, P*(*x, y*) → *αΦ* (*Γ*) →1 →*γ*. Thus, there must be at least one *x* such that *P* = 0 whenever

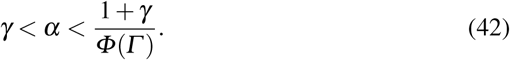

Given that *γ* and *Γ* are small, this condition is satisfied for all but the most extreme values of *α*, corresponding to cases where the assimilation rate of one of the resources is insufficient to replace the C or N of lost biomass.

To demonstrate uniqueness, we differentiate the first of (41) to obtain

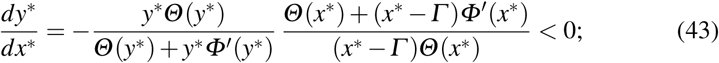

then differentiation of (40) yields

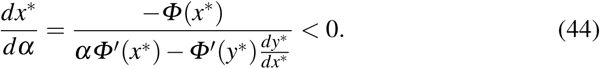

These latter results show that each *α* yields a unique *x*\*and each *x*\*a unique *y*\*; hence, *u*\*is unique for any given *α* in the range given by (42).

#### C.4.2 Stability of u*

To determine stability of *u*\*, we can think of the system (41) as defining *x*\*(*u*\*) and *y*\*(*x*\*(*u*\*)); hence, the stability criterion is

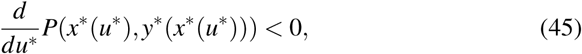

where *P* is given in (23). Differentiating *P*(*x*(*u*), *y*(*x*(*u*)) yields

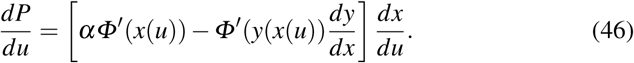

The quantity inside the brackets is always positive, so the stability criterion reduces to *dx*\**/du*\*< 0. Differentiating the second equation of (41) and using (43) yields

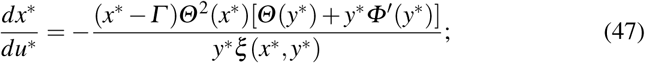

hence, the sign of *dx*\**/du*\*depends on the sign of *ξ*(*x*\*, *y*\*), with stability occurring when *ξ*> 0 or, equivalently,

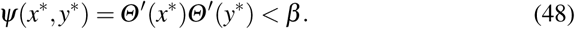

This condition is always satisfied when the SU equations have a unique solution and is sometimes satisfied even when this is not the case.

Note that the combination of the stability criterion *dx*\**/du*\*< 0 with the previous result *dx*\**/dα* < 0 means that *du*\**/dα* > 0 is an equivalent criterion for stability of the equilibrium assimilation ratio. This is an interesting result, given that *u* = *αS/R*. The property *dx*\**/dα* < 0 means that the shoot-root ratio decreases as *α* increases; however, instability occurs only when the decrease in shoot-root ratio with *α* is sufficiently large to overcome the increase in the factor *α*.

### C.5 Details for the PCSU and the kSU

The PCSU has

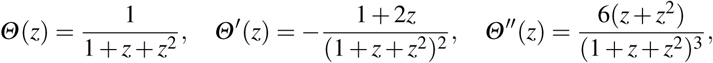

leading to (see Section 4.1)

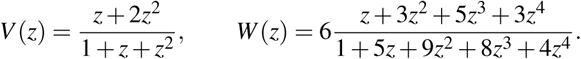

The uniqueness condition (see Section 4.1)

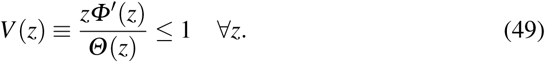

is only satisfied for *z* ≤1, so uniqueness of solutions depends on having *β* ≥*β*_*c*_, which is given asymptotically from (39), with *y*_0_ = 1, *Φ*_1_ = − 1*/*3, and *Φ*_2_ = −4*/*3, as

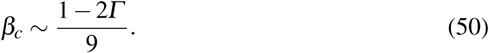

The kSU

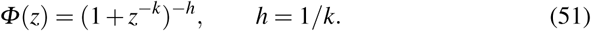

has

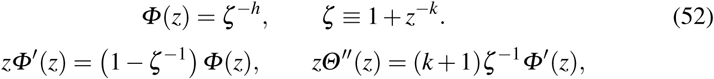

leading to

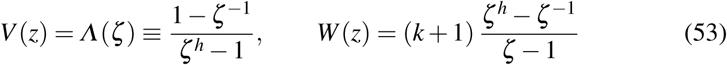

The function *V* is monotone increasing, with *V ∼ k* as *z* →∞, so (49) is satisfied for *k*≤ 1; thus, the SU solution is always unique for that case. If *k* > 1, then *W* is monotone and *β*_*c*_ can be found from (39). The equation *V* (*y*_0_) = 1 can be ultimately rewritten as

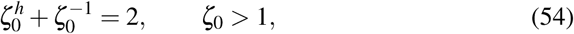

where

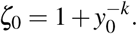

Given 0 < *h* < 1, (54) has a unique solution with *ζ* > 1, yielding a unique solution for *y*_0_. Once *y*_0_ is known, it can be used to calculate

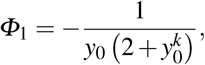

whence

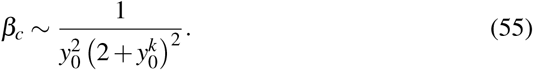

## 1 Appendix D: Optimal Growth with a Symmetric Ratio-Based SU and Global Control

The optimality theory of Iwasa & Roughgarden (1984) and Velten & Richter (1995) does not apply to systems with multiple resources. However, we can address the question of what fixed value of the assimilation ratio *u* leads to the highest rate of balanced growth, and we can run simulations to develop some insight into optimal allocation during the approach to balanced growth.

### D.1 Modification of the model to include resource diversion

Global control can be thought of as a set of allocation rules that replace the default rule in which all C goes directly to the shoot SU and all N available to the root goes to the root SU. A simple way to incorporate global control into our basic model is to assume that a fraction of the input streams can be diverted to the partner without first going to the local SU.

We consider diversion of C and diversion from N separately. Suppose a fraction 0 ≤ *ε* ≤ 1 of the C input in the shoot is shunted directly to the root without having to be rejected by the shoot SU. This changes the C input rate to the shoot SU from *α*_*C*_*S* to *κα*_*C*_*S*, where *κ* = 1 − *ε* serves as an allocation parameter representing the fraction of available C that is sent directly to the shoot SU; thus, *κ* = 1 in the absence of global control. Note that diverting a portion of the C directly to the root does not change the formula for the total C availability because the diverted stream and the rejection flux from the shoot SU are combined. Strategic diversion can still increase the C availability to the root by decreasing *Q*_*S*_.

The remainder of the development of the model equations is largely unchanged by the extra factor of *κ*. Using the same definitions for *x* and *y* (18), the equations for *Q*_*R*_ and *Q*_*S*_ (19) become

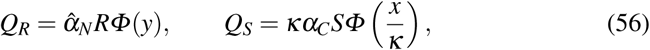

leading to the SU equation (replacing (41))

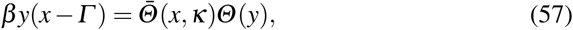

Where

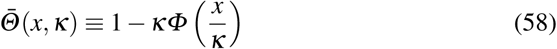

### D.2 Optimal balanced growth rate

Balanced growth means that there is a long-term growth rate *λ* for which

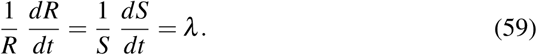

This can occur only at an equilibrium assimilation ratio *u*\*(*κ*), which is associated with the point (*x*\*(*κ*), *y*\*(*κ*)) that satisfies the equation (generalized from (40))

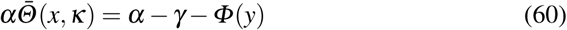

as well as the SU equation (57). From (59), we can identify the growth rate as

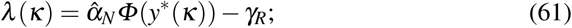

because *Φ* is monotone increasing, the maximum growth rate is achieved at the value of *κ* ∈ [0, 1] for which *y*\*(*κ*) is a maximum. Hence, we may address the issue of optimal *κ* for balanced growth by examining the derivative *dy/dκ*.

Differentiating (60) yields

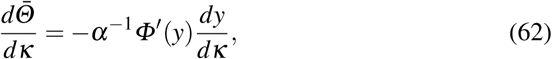

and then (57) yields

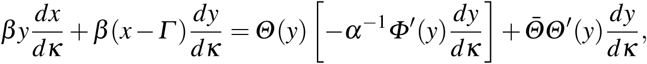

which can be rewritten as

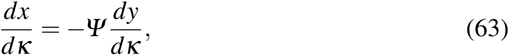

where

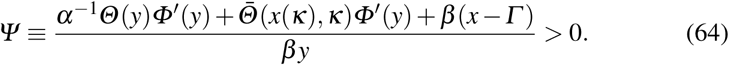

From (58) and (63), we have

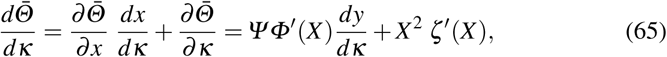

where

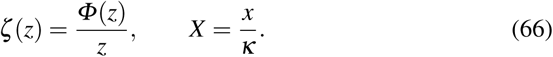

Combining (62) and (65) yields the result

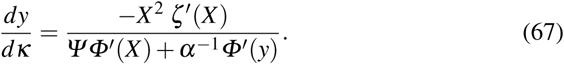

Thus, the sign of *dy/dκ* depends on the sign of *ζ*^′^; in particular, the result shows that diversion of C from the shoot SU always decreases the balanced growth rate whenever the SU function *Φ* satisfies the requirement

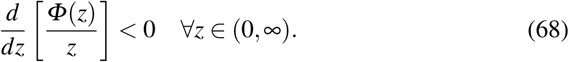

This condition is satisfied by the PCSU, where

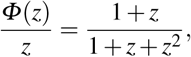

and the kSU, where

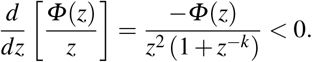

If we instead allow a fraction *ε* of the N available in the root to be diverted directly to the shoot SU, we obtain the requirement that *κ* = 1 − *ε* should be chosen to maximize *x* rather than *y* and obtain a formula analogous to (67) for *dx/dκ*, leading to the same sufficient condition (68) for optimality of *κ* = 1. The more complicated setting in which portions of both resources can be diverted to the partner shows the same result; hence, (68) is a sufficient condition for local allocation to yield optimal balanced growth whenever the equilibrium assimilation ratio is stable.

## Footnotes

1 This omits the resorbed N in the shoots. Functionally, there is no difference between N collected from the environment by roots and N resorbed from senesced roots, since both streams are sent to the root SU. However, N resorbed in the shoot is best omitted from the assimilation ratio, except as noted below.

